# A sensitive period for the development of episodic-like memory in mice

**DOI:** 10.1101/2024.11.06.622296

**Authors:** Adam I Ramsaran, Silvia Ventura, Julia Gallucci, Mitchell L De Snoo, Sheena A Josselyn, Paul W Frankland

## Abstract

Episodic-like memory is a later-developing cognitive function supported by the hippocampus. In mice, the formation of extracellular perineuronal nets in subfield CA1 of the dorsal hippocampus controls the emergence of episodic-like memory during the fourth postnatal week (Ramsaran et al., 2023). Whether the timing of episodic-like memory onset is hard-wired, or flexibly set by early-life experiences during a critical or sensitive period for hippocampal maturation, is unknown. Here, we show that the trajectories for episodic-like memory development vary for mice given different sets of experiences spanning the second and third postnatal weeks. Specifically, episodic-like memory precision developed later in mice that experienced early-life adversity, while it developed earlier in mice that experienced early-life enrichment. Moreover, we demonstrate that early-life experiences set the timing of episodic-like memory development by modulating the pace of perineuronal net formation in dorsal CA1. These results indicate that the hippocampus undergoes a sensitive period during which early-life experiences determine the timing for episodic-like memory development.

## INTRODUCTION

In developing sensory systems, circuit refinement occurs during defined windows of heightened brain plasticity known as critical periods (Hensch, 2005). Rather than occurring spontaneously, functional and then structural refinement of sensory circuits, along with associated developmental shifts in sensory processing, depend on natural sensory experiences occurring during critical periods. For example, depriving the retina of visual stimulation during the period for ocular dominance plasticity in primary visual cortex (V1) may result in lifelong loss of visual acuity (amblyopia), especially if binocular vision is not restored before the closure of the critical period for ocular dominance (Birch, 2013; Chapman & Stryker, 1993; Daw et al., 1992; Espinosa & Stryker, 2012; Holmes et al., 2011; Iny et al., 2006; Morishita & Hensch, 2008). Similarly, impairments in auditory (Harrison et al., 2005; Mowery et al., 2015; Polley et al., 2013; Villers-Sidani et al., 2007; Zhang et al., 2002) and somatosensory (Erzurumlu & Gaspar, 2012; Fox, 1992; McRae et al., 2007) processing can result from sensory deprivation during early postnatal development, underscoring the critical role of early-life experiences in shaping neural circuits across the brain.

Outside of sensory systems, postnatal experience has long been recognized to influence social, emotional, and cognitive development across species (Birnie & Baram, 2022; Freedman et al., 1961; Hebb, 1947; Lorenz, 1935; Opendak et al., 2017; Scott, 1962). With respect to cognition, adverse or enriching experiences occurring during infancy or childhood in humans (Amso et al., 2019; Decker et al., 2020; Evans & Schamberg, 2009; Lambert et al., 2017) and rodent model species (Ivy et al., 2020; Malave et al., 2022; Molet et al., 2016; Woodcock & Richardson, 2000) can have profound effects on mnemonic function later in life, despite the memories for these experiences being long-forgotten (Akers et al., 2014; Madsen & Kim, 2016; Ramsaran et al., 2019; Rubin & Schulkind, 1997). These findings are consistent with recent proposals that the hippocampus undergoes a critical period during which it becomes competent to form episodic memories (Alberini & Travaglia, 2017) and predict that childhood experiences modulate the trajectory of hippocampal maturation in a manner similar to experience-dependent maturation of sensory cortices. Yet, the specific neurobiological mechanisms in the hippocampus that mediate the effects of experience on memory development have not been identified.

Recently, we reported that the capacity of the hippocampus to form precise episodic-like memories emerges between postnatal days 20 (P20) and 24 in mice (Ramsaran et al., 2023). In the CA1 region of the hippocampus of juvenile (≤P20) mice, events are allocated to dense neuronal engram ensembles, whereas in the CA1 of older (≥P24) mice, the same events are allocated to sparse neuronal engram ensembles. We found that the onset of sparse engram formation and episodic-like memory during the fourth postnatal week requires a developmental increase in circuit inhibition mediated by postnatal accumulation of extracellular perineuronal nets (PNNs) – a canonical molecular mechanism for terminating critical periods in sensory cortices (Carulli et al., 2010; Carulli & Verhaagen, 2021; Pizzorusso et al., 2002; D. Wang & Fawcett, 2012). Because cortical PNN formation is triggered by sensory experience (McRae et al., 2007; Ye & Miao, 2013), here we addressed whether early postnatal experiences affect the timing of hippocampal PNN formation and the emergence of episodic-like memory function. We found that early-life adversity or enrichment respectively decelerate or accelerate the development of sparse engrams and episodic-like memory precision by modulating the pace of PNN maturation in CA1 of the dorsal hippocampus. Thus, the timing of episodic-like memory onset is not fixed, but rather determined by experiences occurring during infancy and childhood, which set the pace for hippocampal maturation.

## RESULTS

### Experience-dependent PNN formation as a brain-wide mechanism for neural circuit maturation

In many parts of the brain, accumulation of PNNs marks the end of critical periods for neural and/or behavioral maturation (Gogolla et al., 2009; Mirzadeh et al., 2019; Pizzorusso et al., 2002). In sensory cortices of rodents, where PNNs are essential for both mature parvalbumin (PV) interneuron function and sensory processing, trajectories for PNN formation vary significantly, but reliably coincide with the termination of region-specific critical periods. To demonstrate this principle, we used *Wisteria floribunda* agglutinin (WFA) labeling to examine the developmental trajectories for PNN formation in two sensory cortical regions with well-described, non-overlapping critical periods: layer IV of both V1 and the barrel fields of primary somatosensory cortex (S1BF). Consistent with the termination of ocular dominance plasticity during the fifth postnatal week (Gordon & Stryker, 1996) and barrel field plasticity before the second postnatal week (Fox, 1992), we observed a slower accumulation of WFA^+^ PNNs surrounding PV^+^ interneurons in V1 beginning one week after eye opening, which contrasted with an abundance of WFA^+^ PNNs in S1BF observed as early as P12 in the same mice (**Figure 1A-B**). Therefore, it is possible to track the decline of critical period plasticity by examining the local developmental trajectory for PNN formation.

**Figure 1.**
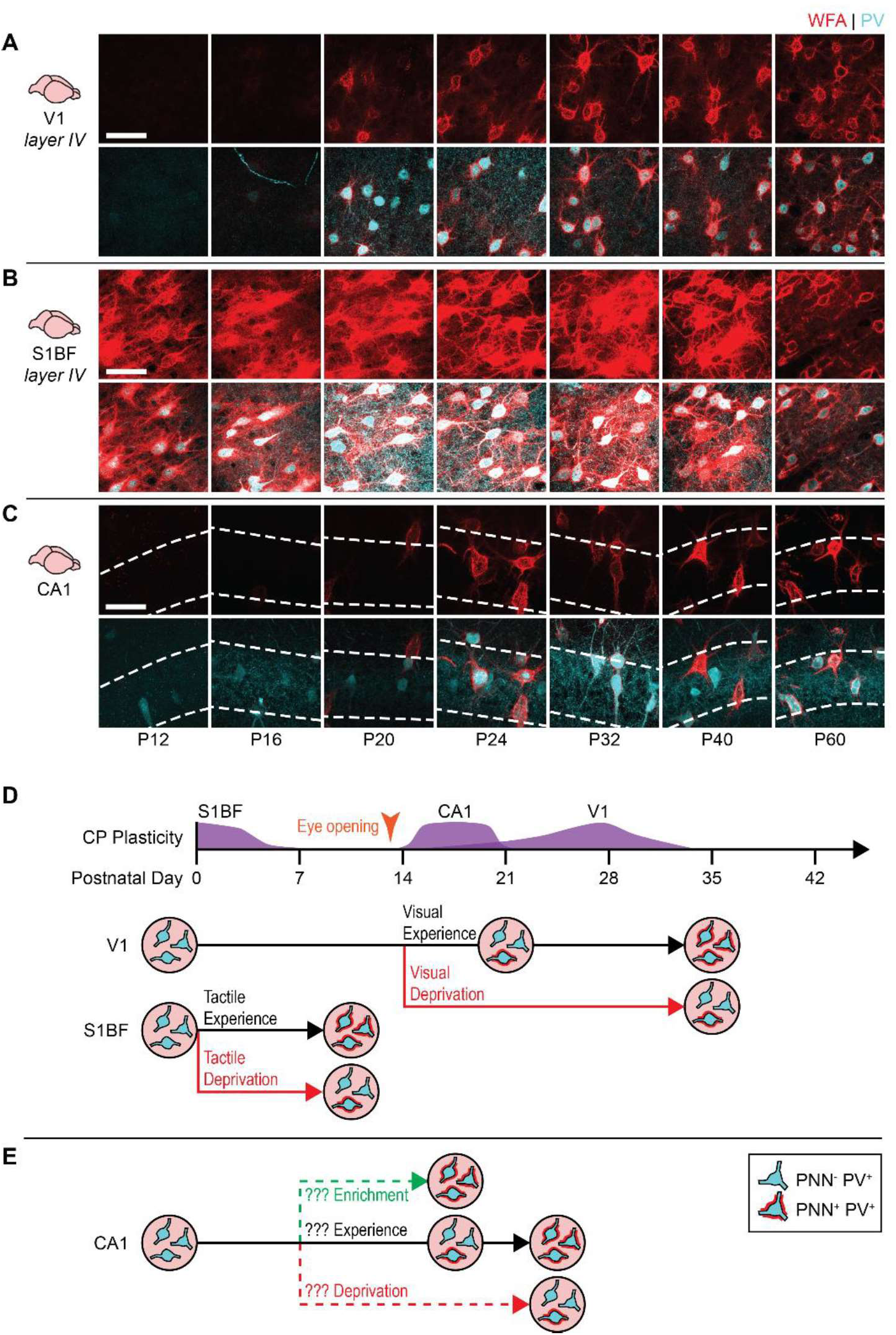
Experience-dependent critical period mechanisms may control the maturation of episodic-like memory. (A) In primary visual cortex (V1), the formation of WFA^+^ PNNs around PV^+^ interneurons begins after eye opening and is largely completed by the fifth postnatal week, consistent with the timing of critical period closure for ocular dominance plasticity (Gordon & Stryker, 1996). (B) In the barrel fields of primary somatosensory cortex (S1BF), the formation of WFA^+^ PNNs around PV^+^ interneurons has already occurred by the second postnatal week, consistent with the timing of critical period closure for barrel field plasticity (Fox, 1992). (C) In the dorsal CA1 of the hippocampus, the formation of WFA^+^ PNNs around PV^+^ interneurons occurs rapidly during the fourth postnatal week. (D) PNN-dependent critical period closure requires specific postnatal experiences during the critical period. In mice, vision loss or dark rearing from birth impairs PNN formation in the binocular zone of primary visual cortex (V1), resulting in impaired visual acuity (Hensch, 2005; Huang et al., 1999; Pizzorusso et al., 2002; Ye & Miao, 2013). Likewise, whisker removal from birth precludes PNN formation in layer 4 of barrel cortex (Fox, 1992; McRae et al., 2007). (E) If PNN formation in the dorsal CA1 marks the end of a hippocampal critical period, deprivation or enrichment should shift the developmental trajectories for PNN formation and episodic-like memory. Scale bar, 50 µm.

We next applied this principle to the CA1 subfield of the dorsal hippocampus. In a previous study, we showed that PNN formation in this region is responsible for the developmental onset of hippocampus-dependent memory precision, which is a core feature of episodic-like memory (Ramsaran et al., 2019, 2023). As reported before, we found that WFA^+^ PNN formation around PV^+^ interneurons occurred rapidly between P20 and P24 (**Figure 1C**), tracking the emergence of hippocampal memory precision in rodents (Anderson & Riccio, 2005; Ramsaran et al., 2023). These results suggest that PNN formation in CA1 and sensory cortices play similar roles in the maturation of hippocampal memory and sensory processing, respectively, and may indicate the existence of a critical period for episodic-like memory development in the hippocampus.

Postnatal experiences are essential for neural and behavioral maturation during critical periods. With respect to sensory critical periods, these experiences must be both modality-specific and timed to specific postnatal epochs. For example, in rodents, PNN formation in the binocular zone of V1 (Pizzorusso et al., 2002; Ye & Miao, 2013) and S1BF (Nowicka et al., 2009) require visual and somatosensory input to the retina and whiskers, respectively, and can be impaired by sensory deprivation that spans each region’s defined critical period (Carulli et al., 2010; McRae et al., 2007; Pizzorusso et al., 2002; Ye & Miao, 2013) (**Figure 1D**). To determine whether postnatal formation of PNNs in the CA1 subfield of the dorsal hippocampus is analogously regulated by experience, we manipulated the early rearing environment, which modulates neuronal activity in the immature rodent hippocampus (Pompeiano & Colonnese, 2023; Xue et al., 2024), rather than specific sensory experiences. To this end, we used experiential interventions in the weeks preceding typical CA1 PNN formation (i.e., P6 to P18-19), predicting that, like for sensory systems, the timing of hippocampal memory maturation is regulated in an experience-dependent manner (**Figure 1E**).

### Early-life adversity decelerates, but does not abolish, the maturation of CA1 PNNs and episodic-like memory

We first asked whether deprivation during postnatal development would impair the formation of dorsal CA1 PNNs and the maturation of episodic-like memory. Different forms of adversity during early life (early-life adversity, ELA) negatively affect neural and behavioral aspects of hippocampal development (Elliott & Richardson, 2019; Huot et al., 2002; Murthy et al., 2019; Raineki et al., 2019), but the impact of ELA on dorsal CA1 PNNs is poorly understood. Therefore, we subjected mice to an ELA protocol (Murthy et al., 2019) consisting of daily maternal separation (6 h/day from P6 to P16, inclusive) followed by early weaning (on P17, compared to typical weaning on P21; **Figure 2A**). We then examined PNNs and memory precision on P24, when episodic-like memory has recently matured. In mice experiencing ELA, weight gain was modestly reduced compared to control mice during and after the maternal separation period (**Figure S1A**); however, the timing of eye opening — a developmental milestone gating shifts in critical period plasticity across multiple sensory systems (Mowery et al., 2016; Smith & Trachtenberg, 2007) — was unaffected (**Figure S1B**). Therefore, potential changes in hippocampal plasticity following ELA cannot be explained by differences in the onset of vision. In P24 mice with a history of ELA, hippocampal PNN formation was attenuated, as we observed reduced amounts of WFA^+^ PNNs and PV^+^ interneurons in CA1 compared to control mice (**Figures 2B-C**). By contrast, CA3 PNNs, which condense around PV^+^ interneurons >1 week earlier than CA1 PNNs (Ramsaran et al., 2023), were unaffected by the ELA intervention (**Figure 2D-E**), suggesting that a different, likely earlier, sensitive period for experience-dependent PNN formation in CA3.

**Figure 2.**
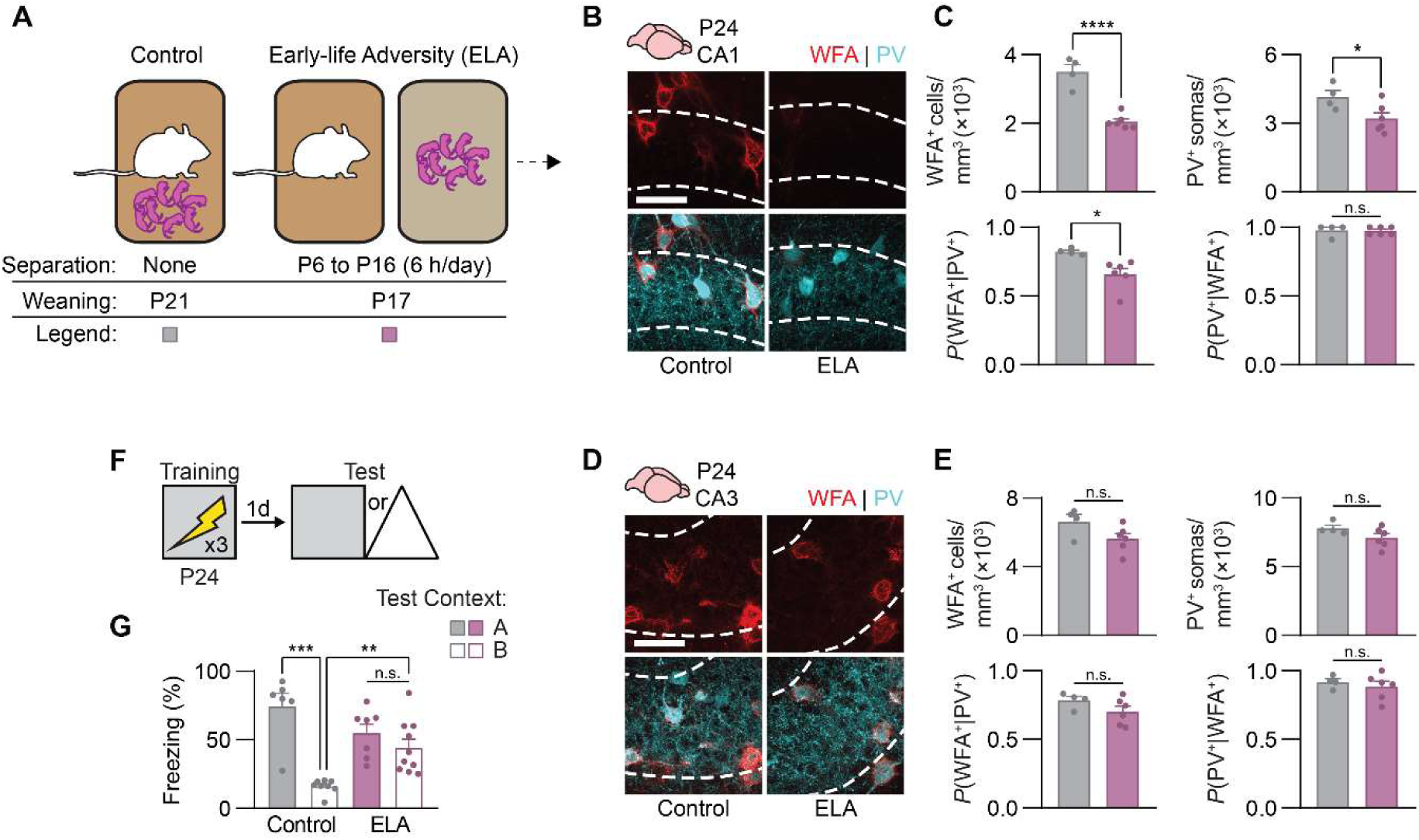
Early-life adversity impairs the maturation of perineuronal nets in CA1 and memory precision in juvenile mice. (A) Mice experienced adversity in the forms of daily maternal separation from P6 to P16 followed by early weaning on P17 (ELA group), or no separation and conventional weaning (Control group). (B) Labeling of WFA^+^ PNNs (red) and PV^+^ interneurons (cyan) in dorsal CA1 of Control and ELA mice on P24. (C) P24 mice with a history of adversity had fewer WFA^+^ PNNs (unpaired *t*-test, *t*_8_ = 7.17, *P* < 0.0001), PV^+^ interneurons (unpaired *t*-test, *t*_8_ = 2.47), and PNN-enwrapped PV^+^ interneurons (unpaired *t*-test, *t*_8_ = 2.99, *P* < 0.05), but not PV^-^ PNNs (unpaired *t*-test, *t*_8_ = 0.11, *P* = 0.91) in dorsal CA1, compared with Control mice. (D) Labeling of WFA^+^ PNNs (red) and PV^+^ interneurons (cyan) in dorsal CA3 of Control and ELA mice on P24. (E) There was no difference in WFA^+^ PNNs (unpaired *t*-test, *t*_8_ = 2.00, *P* = 0.081), PV^+^ interneurons (unpaired *t*-test, *t*_8_ = 1.65, *P* = 0.13), PNN-enwrapped PV^+^ interneurons (unpaired *t*-test, *t*_8_ = 1.50, *P* = 0.17), or PV^-^ PNNs (unpaired *t*-test, *t*_8_ = 0.67, *P* = 0.52) in dorsal CA3 of P24 Control and ELA mice. (F) Experimental schedule for assessing memory precision. Mice were trained in a contextual fear conditioning task on P24, and 1 d later, they were tested in the same context (Context A) or a similar, novel context (Context B). (G) P24 mice with a history of adversity showed imprecise contextual fear memories, compared with Control mice that showed age-appropriate memory precision (ANOVA, experience × test context interaction: *F*_1,27_ = 14.27, *P* < 0.001). Data points represent individual mice with mean ± SEM. Scale bar, 50 µm. \**P* < 0.05, \*\**P* < 0.01, \*\*\**P* < 0.001, \*\*\*\**P* < 0.0001.

Given the impairment in CA1 PNN formation in ELA mice on P24, we predicted that these mice would show imprecise memory formation comparable to younger (P20) mice. To test this, separate groups of mice experienced ELA or control rearing. The mice were then trained in contextual fear conditioning on P24, and memory precision was assessed the following day by testing the mice in the same apparatus (Context A) or a novel, but similar apparatus (Context B) (**Figure 2F**). We used freezing behavior as a proxy for memory recall. While mice that did not experience ELA showed age-appropriate memory precision during the test, freezing more in Context A than Context B, mice previously experiencing ELA showed imprecise memories, freezing equally in Contexts A and B (**Figure 2G**). In contrast to previous studies reporting sex-dependent effects of ELA on behavioral and extrahippocampal PNN development (Gildawie et al., 2020; Goodwill et al., 2019; Guadagno et al., 2020), we found that ELA-induced impairments in the development of CA1 PNNs and memory precision did not differ between sexes (**Figure S2A-C**). Together, these data suggest that ELA impairs both neural and behavioral maturation of episodic-like memory.

If the hippocampus undergoes a critical period for episodic-like memory, ELA should completely block the emergence of PNN-dependent memory precision. Alternatively, ELA may delay the development of PNN-dependent memory precision, suggesting that there is a sensitive (but not critical) period for episodic-like memory development (Hensch, 2005). To test these possibilities, we repeated the ELA intervention and examined CA1 PNNs and memory precision in young adult mice on P60 (**Figure 3A**). In dorsal CA1, WFA^+^ PNNs and PV^+^ interneurons did not differ between adult mice with and without a history of ELA (**Figure 3B-C**). Consistent with having formed PNNs in CA1 sometime between juvenility and young adulthood, memory precision was intact in P60 ELA mice (**Figure 3D-E**). Despite demonstrating normal memory precision, P60 ELA mice displayed an anxiety-like behavioral phenotype in the open field (**Figure 3F**), consistent with earlier work (Qin et al., 2021; X.-D. Wang et al., 2012). Therefore, the affective behavioral outcomes of ELA are more enduring than the cognitive outcomes, with ELA delaying, but not abolishing, the maturation of episodic-like memory. These data support the idea that the hippocampus undergoes a sensitive period for episodic-like memory development.

**Figure 3.**
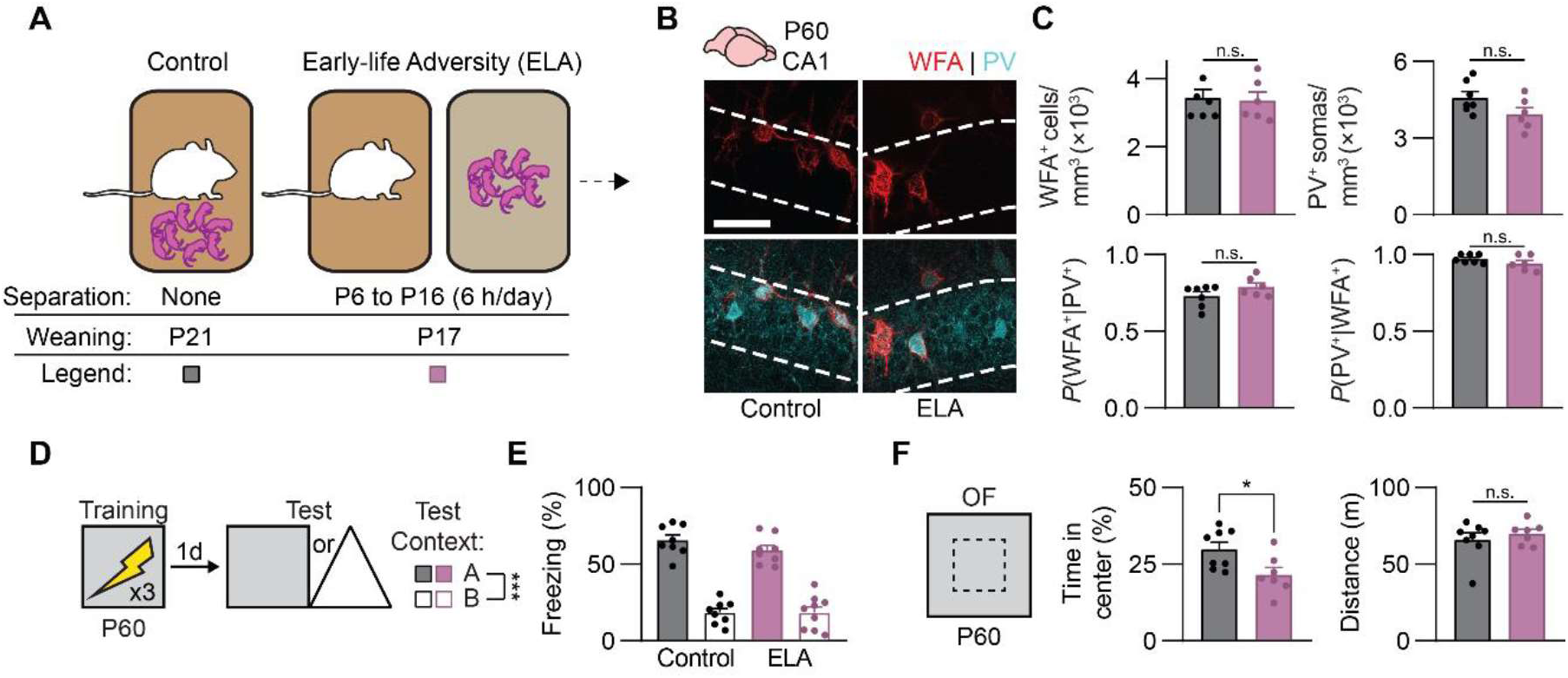
Early-life adversity does not block the maturation of perineuronal nets in CA1 and memory precision in adult mice. (A) Mice experienced adversity in the forms of daily maternal separation from P6 to P16 followed by early weaning on P17 (ELA group), or no separation and conventional weaning (Control group). (B) Labeling of WFA^+^ PNNs (red) and PV^+^ interneurons (cyan) in dorsal CA1 of Control and ELA mice on P60. (C) There was no difference in WFA^+^ PNNs (unpaired *t*-test, *t*_11_ = 0.25, *P* = 0.81), PV^+^ interneurons (unpaired *t*-test, *t*_11_ = 1.88, *P* = 0.088), PNN-enwrapped PV^+^ interneurons (unpaired *t*-test, *t*_11_ = 1.58, *P* = 0.14), or PV^-^ PNNs (unpaired *t*-test, *t*_11_ = 1.38, *P* = 0.19) in dorsal CA1 of P60 Control and ELA mice. (D) Experimental schedule for assessing memory precision. Mice were trained in a contextual fear conditioning task on P60, and 1 d later, they were tested in the same context (Context A) or a similar, novel context (Context B). (E) P60 Control and ELA mice showed adult-like memory precision (ANOVA, main effect of test context: *F*_1,29_ = 171.78, *P* < 0.0001). (F) Compared to Control mice, ELA mice on P60 showed increased anxiety-like behavior (unpaired *t*-test, *t*_13_ = 2.53, *P* < 0.05) but similar levels of locomotion (unpaired *t*-test, *t*_13_ = 0.72, *P* = 0.48) in the open field. Data points represent individual mice with mean ± SEM. Scale bar, 50 µm. \**P* < 0.05, \*\**P* < 0.01, \*\*\**P* < 0.001, \*\*\*\**P* < 0.0001.

### BDNF treatment following early-life adversity restores CA1 PNNs and rescues episodic-like memory

We hypothesized that the delayed onset of episodic-like memory precision in ELA mice was mediated specifically by the later formation of dorsal CA1 PNNs following ELA. However, dysregulation of neural circuit maturation by ELA is not restricted to the hippocampus. To causally demonstrate that ELA delays memory precision by delaying maturation of CA1 PNNs (i.e., to rule out other effects of ELA on brain maturation), we sought to “rescue” the impairment in memory precision in ELA mice using a CA1 - specific treatment. We decided to use brain-derived neurotrophic factor (BDNF) as a treatment following ELA because BDNF serves as a key regulator of critical period plasticity and neural circuit maturation (Hanover et al., 1999; Hensch, 2005; Huang et al., 1999; Travaglia et al., 2016). Via molecular cascades initiated by binding its receptor, tyrosine receptor kinase B (TrkB), or via direct binding to chondroitin-sulfate proteoglycans (CSPGs) (Fawcett et al., 2019), BDNF treatment induces CSPG expression in neurons (Willis et al., 2021), stimulates precocial PNN formation in the juvenile mouse hippocampus (Ramsaran et al., 2023), improves hippocampus-dependent memory precision in adult rats (Miranda et al., 2018), and rescues impairments in cortical function caused by sensory deprivation (Gianfranceschi et al., 2003).

To determine whether BDNF treatment would restore age-appropriate PNN density and memory following ELA, on P21 we microinfused solution containing recombinant BDNF, or vehicle, into dorsal CA1 of separate groups of mice (**Figure 4A**). As shown above, WFA^+^ PNN formation around PV^+^ interneurons was attenuated in P24 mice that experienced ELA and that were treated with vehicle. By contrast, BDNF treatment on P21 restored WFA^+^ PNNs to age-appropriate levels in ELA mice by P24 (**Figure 4B-C**). Moreover, in separate groups of ELA mice, we observed age-appropriate memory precision in P24 mice that received BDNF infusions into CA1 days before contextual fear conditioning, compared with mice that received vehicle infusions (**Figure 4D-E**). These experiments indicate that ELA, a form of deprivation, shifts the trajectory of episodic-like memory development via its effects on hippocampal PNN formation. Similar to V1 (Gianfranceschi et al., 2003), restoring the local maturational state of CA1 following deprivation rescues age-appropriate neural circuit function.

**Figure 4.**
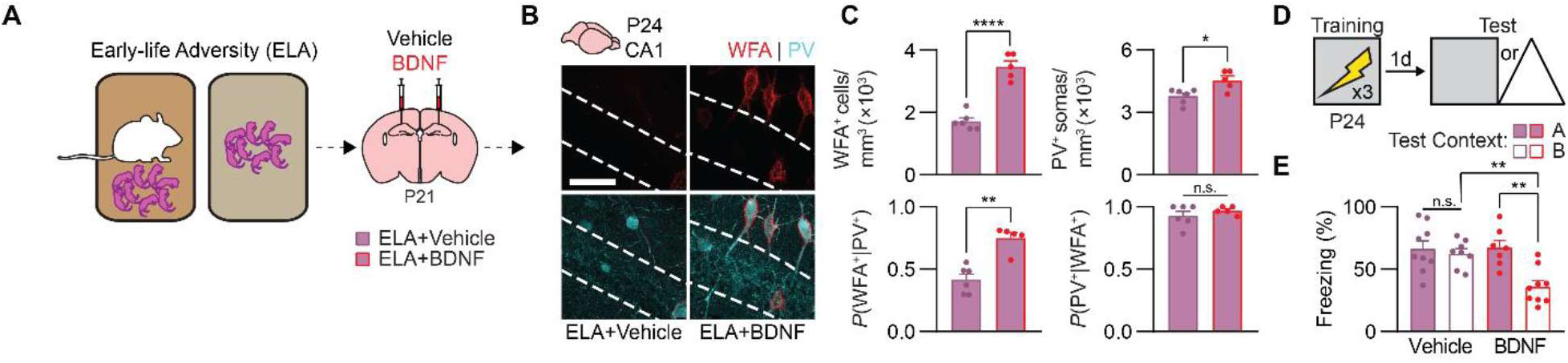
BDNF treatment restores perineuronal nets in CA1 and memory precision in juvenile mice raised in adverse conditions. (A) Mice experienced adversity followed by infusions of Vehicle or BDNF into CA1 on P21. (B) Labeling of WFA^+^ PNNs (red) and PV^+^ interneurons (cyan) in dorsal CA1 of ELA mice treated with Vehicle or BDNF. (C) P24 mice treated with BDNF following early-life adversity had more WFA^+^ PNNs (unpaired *t*-test, *t*_9_ = 8.46, *P* < 0.0001), PV^+^ interneurons (unpaired *t*-test, *t*_9_ = 3.01, *P* < 0.05), and PNN-enwrapped PV^+^ interneurons (unpaired *t*-test, *t*_9_ = 5.25, *P* < 0.001), but not more PV^-^ PNNs (unpaired *t*-test, *t*_9_ = 0.97, *P* = 0.36), than mice treated with Vehicle. (D) Experimental schedule for assessing memory precision. Mice were trained in a contextual fear conditioning task on P24, and 1 d later, they were tested in the same context (Context A) or a similar, novel context (Context B). (E) P24 mice treated with BDNF following early-life adversity showed age-appropriate contextual fear memory precision, compared with P24 mice treated with Vehicle, that showed imprecise memories (ANOVA, treatment × test context interaction: *F*_1,29_ = 6.55, *P* < 0.05). Data points represent individual mice with mean ± SEM. Scale bar, 50 µm. \**P* < 0.05, \*\**P* < 0.01, \*\*\**P* < 0.001, \*\*\*\**P* < 0.0001.

### Early-life enrichment accelerates the maturation of CA1 PNNs and episodic-like memory

That ELA, a form of deprivation, decelerates the maturation of hippocampus-dependent episodic-like memory suggests that enriching experiences during early postnatal development might have the opposite effect. Early-life environmental enrichment (ELE) enhances both physical (i.e., object and spatial novelty) and social (i.e., maternal behavior) aspects of pups’ postnatal experience and results in precocial maturation of V1-dependent visual acuity (Cancedda et al., 2004; Sale et al., 2004). Additionally, prolonged environmental enrichment improves hippocampus-dependent memory across different stages of the rodent lifespan (Duffy et al., 2001; Frick et al., 2003), although the effects of perinatal enrichment on memory in juvenile rodents has not been characterized.

To test whether ELE promotes hippocampal memory maturation in developing mice, we continuously housed mouse pups (and their mother) in large cages containing multiple pieces of physical enrichment (i.e., domes, tubes, etc.) and extra nesting materials. Enrichment began on P6 and ended on P19, when the mice were returned to standard housing conditions (**Figure 5A**). The physical enrichment was replaced or rearranged every few days to ensure that the environment remained novel throughout the duration of the intervention. By contrast, control mice remained in standard housing throughout the experiment. ELE did not affect weight gain or eye opening in developing mice (**Figure S1A-B**). We then examined PNNs in the dorsal hippocampus on P20; a time point when PNNs in normally-reared mice have not yet finished forming. We observed precocial accumulation of WFA^+^ PNNs around PV^+^ interneurons in dorsal CA1 of P20 mice that experienced ELE, compared to P20 control mice (**Figures 5B-C**). No differences in WFA^+^ PNNs or PV^+^ interneurons were observed between ELE and control mice in dorsal CA3 (**Figure 5D-E**), again suggesting that dorsal CA1 PNN maturation is more sensitive to postnatal experiences spanning the second and third postnatal weeks. In separate groups of mice, we next tested memory precision using contextual fear conditioning. Consistent with the earlier formation of CA1 PNNs, ELE mice showed precise contextual fear memories following training on P20, whereas control mice showed age-appropriate memory imprecision (**Figure 5F-G**). The effects of ELE on CA1 PNN formation and memory precision did not appear to be influenced by sex (**Figure S2D-F**). These experiments indicate that beneficial postnatal experiences, such as those afforded by ELE, accelerate maturation of the hippocampal PNNs and memory precision.

**Figure 5.**
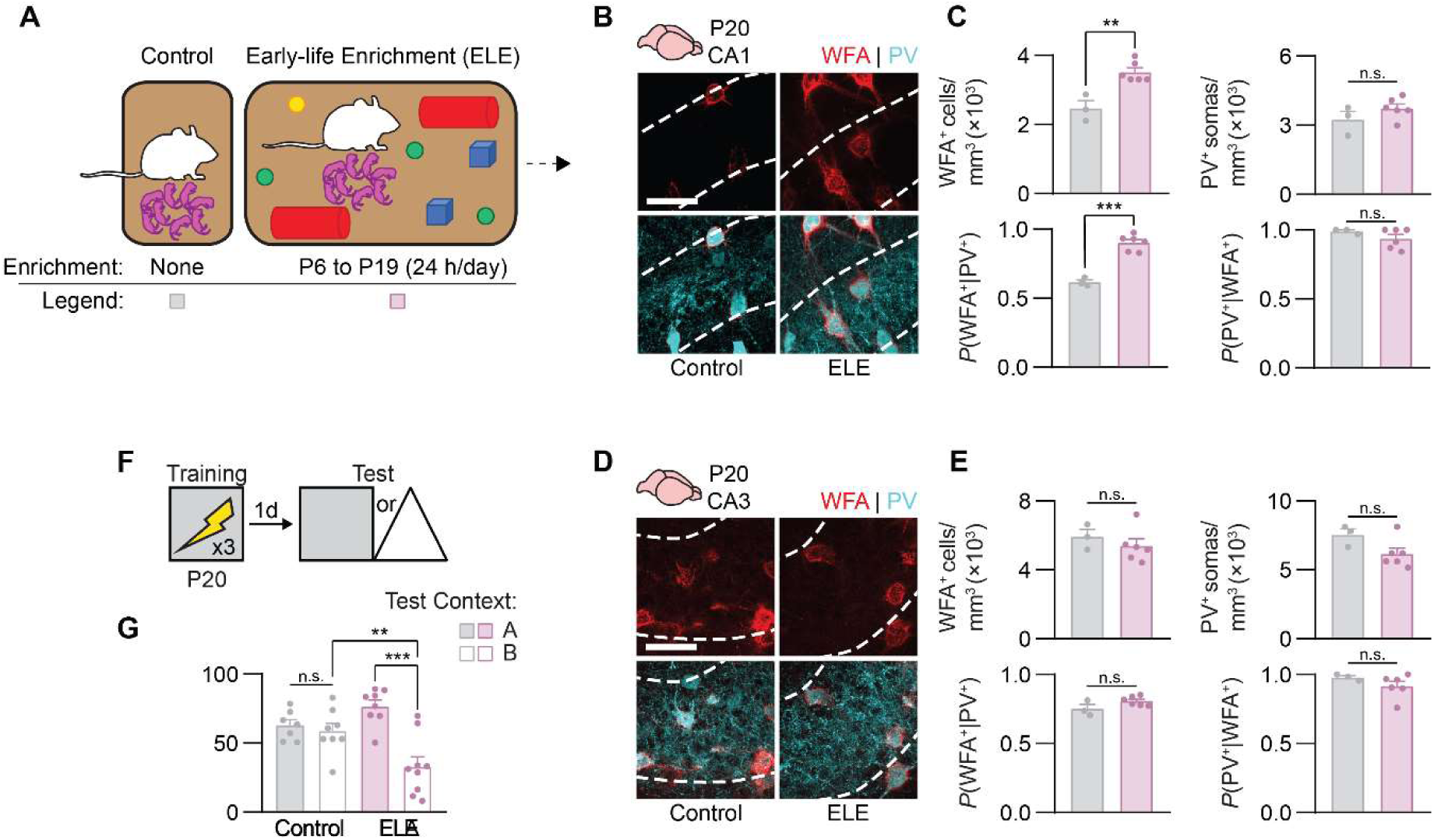
Early-life enrichment induces precocial maturation of perineuronal nets in CA1 and memory precision in preweaning mice. (A) Mice were housed in large cages containing enrichment from P6 to P19 (ELE group) or were housed conventionally (Control group). (B) Labeling of WFA^+^ PNNs (red) and PV^+^ interneurons (cyan) in dorsal CA1 of Control and ELE mice on P20. (C) P20 mice previously raised in enriched environments had more WFA^+^ PNNs (unpaired *t*-test, *t*_7_ = 4.69, *P* < 0.01) and PNN-enwrapped PV^+^ interneurons (unpaired *t*-test, *t*_7_ = 7.84, *P* < 0.001) and equivalent PV^+^ interneurons (unpaired *t*-test, *t*_7_ = 1.38, *P* = 0.21 and PV^-^ PNNs (unpaired *t*-test, *t*_7_ = 1.21, *P* = 0.26) in dorsal CA1, compared with Control mice. (D) Labeling of WFA^+^ PNNs (red) and PV^+^ interneurons (cyan) in dorsal CA3 of Control and ELE mice on P20. (E) There was no difference in WFA^+^ PNNs (unpaired *t*-test, *t*_8_ = 2.00, *P* = 0.081), PV^+^ interneurons (unpaired *t*-test, *t*_8_ = 1.65, *P* = 0.13), PNN-enwrapped PV^+^ interneurons (unpaired *t*-test, *t*_8_ = 1.50, *P* = 0.17), or PV^-^ PNNs (unpaired *t*-test, *t*_8_ = 0.67, *P* = 0.52) in dorsal CA3 of P20 Control and ELA mice. (F) Experimental schedule for assessing memory precision. Mice were trained in a contextual fear conditioning task on P20, and 1 d later, they were tested in the same context (Context A) or a similar, novel context (Context B). (G) P20 mice previously raised in enriched environments showed precise contextual fear memories, compared with Control mice that showed age-appropriate memory imprecision (ANOVA, experience × test context interaction: *F*_1,28_ = 11.64, *P* < 0.01). Data points represent individual mice with mean ± SEM. Scale bar, 50 µm. \**P* < 0.05, \*\**P* < 0.01, \*\*\**P* < 0.001, \*\*\*\**P* < 0.0001.

### ChABC treatment following early-life enrichment digests CA1 PNNs and reverses episodic-like memory enhancement

Environmental enrichment engages multiple neural circuits across the brain (Li et al., 2020), raising the possibility that the precocial onset of episodic-like memory precision seen in P20 ELE mice was supported by extrahippocampal mechanisms. To determine whether the early formation of CA1 PNNs accounted for the precocial onset of memory precision in these mice, we used the enzyme chondroitinase ABC (ChABC) to digest CSPGs found in PNNs (Pizzorusso et al., 2002; Ramsaran et al., 2023). To this end, we generated two groups of mice raised in enriched environments from P6, and, before returning them to standard housing on P18, we microinfused ChABC, or the control enzyme penicillinase, into dorsal CA1 (**Figure 6A**). We examined WFA^+^ PNNs and PV^+^ interneurons in dorsal CA1 on P20, as before. Mice that previously experienced enrichment and that were treated with penicillinase had elevated levels of WFA^+^ PNNs surrounding PV^+^ interneurons in dorsal CA1. By contrast, WFA^+^ PNNs in ELE mice were digested by ChABC (**Figure 6B-C**). Consistent with the reversal of ELE-induced PNN formation in CA1 by ChABC, ChABC treatment in juvenile mice raised in enrichment reinstated memory imprecision typical of juvenile mice raised under control conditions (**Figure 6D-E**). Thus, ELE matures episodic-like memory precision in mice by promoting precocial development of PNNs in dorsal CA1.

**Figure 6.**
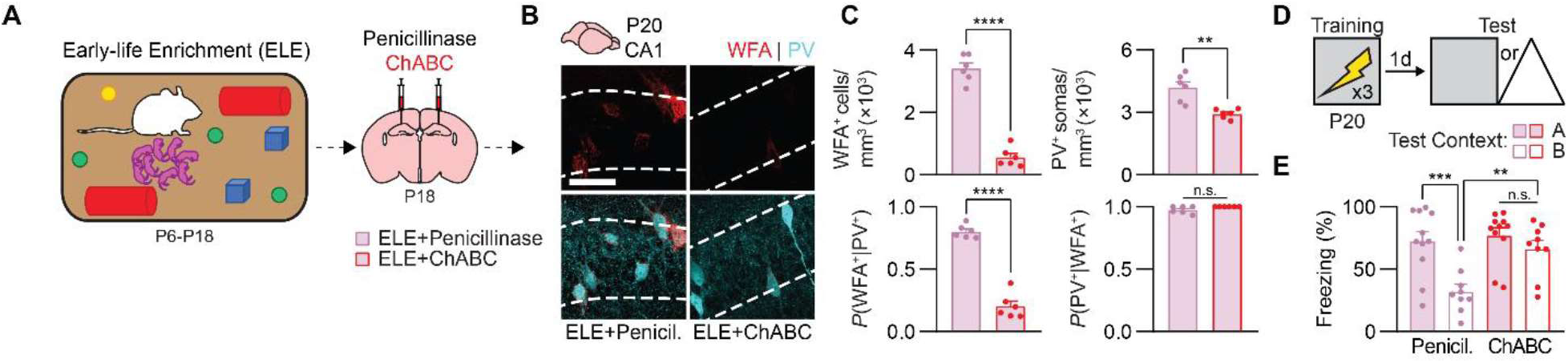
ChABC treatment disrupts perineuronal nets in CA1 and memory precision in preweaning mice raised in enriched conditions. (A) Mice raised in enriched environments from P6 to P18 were infused with Penicillinase or ChABC into CA1 on P18. (B) Labeling of WFA^+^ PNNs (red) and PV^+^ interneurons (cyan) in dorsal CA1 of ELE mice treated with Penicillinase or ChABC. (C) P20 mice treated with ChABC following early-life enrichment had fewer WFA^+^ PNNs (unpaired *t*-test, *t*_10_ = 13.17, *P* < 0.0001), PV^+^ interneurons (unpaired *t*-test, *t*_10_ = 4.49, *P* < 0.01), and PNN-enwrapped PV^+^ interneurons (unpaired *t*-test, *t*_10_ = 12.84, *P* < 0.0001), but not fewer PV^-^ PNNs (unpaired *t*-test, *t*_10_ = 2.08, *P* = 0.06), than mice treated with Penicillinase. (D) Experimental schedule for assessing memory precision. Mice were trained in a contextual fear conditioning task on P20, and 1 d later, they were tested in the same context (Context A) or a similar, novel context (Context B). (E) P20 mice treated with ChABC following early-life enrichment showed age-appropriate contextual fear memory imprecision, compared with P20 mice treated with Penicillinase, that showed precise memories (ANOVA, infusion × test context interaction: *F*_1,36_ = 4.51, *P* < 0.05). Data points represent individual mice with mean ± SEM. Scale bar, 50 µm. \**P* < 0.05, \*\**P* < 0.01, \*\*\**P* < 0.001, \*\*\*\**P* < 0.0001.

### Early-life experience shapes trajectory for neuronal allocation to a sparse engram ensemble via PNNs

Declines or impairments memory precision have been associated with reduced sparsity of memory representations at the level of neuronal engram ensembles (Guayasamin et al., 2024; Lesuis et al., 2021; Poll et al., 2020; Ruediger et al., 2011). Accordingly, we showed that in CA1 of the dorsal hippocampus, PNN-dependent changes in PV^+^ interneuron function control the shift from dense-to-sparse engram formation between P20 and P24, demonstrating causal links between maturation of hippocampal PNNs, neuronal allocation to an engram ensemble, and memory precision (Ramsaran et al., 2023). Given these findings, next we asked whether early-life interventions (ELA and ELE), via their effects on PNN formation, produce changes in memory precision in developing mice by altering how neurons are allocated to engram ensembles in CA1.

To examine how PNN maturational state set by early-life experiences potentially alters the process of neuronal allocation to an engram, we examined c-Fos and PV expression in dorsal CA1 of P20 and P24 mice following encoding a contextual fear memory (**Figure 7A**). To dissociate the effects of early experiences from PNN maturational state in dorsal CA1, we used 6 groups of mice including mice with typically developing CA1 PNNs (P20 Control, P24 Control groups); mice with accelerated or decelerated CA1 PNN development (P20 ELE+Penicillinase, P24 ELA+Vehicle groups); and mice that received the experiential interventions, but whose CA1 PNN maturational state was ‘reset’ using pharmacology before memory encoding (P20 ELE+ChABC, P24 ELA+BDNF groups). As reported previously (Ramsaran et al., 2023), we observed an approximately 2-fold reduction in c-Fos expression following memory encoding between P20 and P24, reflecting an age-dependent increase in neuronal engram ensemble sparsity (**Figure 7B**). In mice that were raised in enriched or adverse environments, c-Fos^+^ engram ensemble sparsity tracked changes in PNN maturational state, rather than age. Specifically, experiential groups that had low levels of PNNs in dorsal CA1 (P20 ELE+ChABC, P24 ELA+Vehicle; **Figures 4B-C, 6B-C**) formed dense engram ensembles compared to groups that had high levels of PNNs in dorsal CA1 (P20 ELE+Penicillinase, P24 ELA+BDNF; **Figures 4B-C, 6B-C**) that formed sparse engram ensembles (**Figure 7B**).

**Figure 7.**
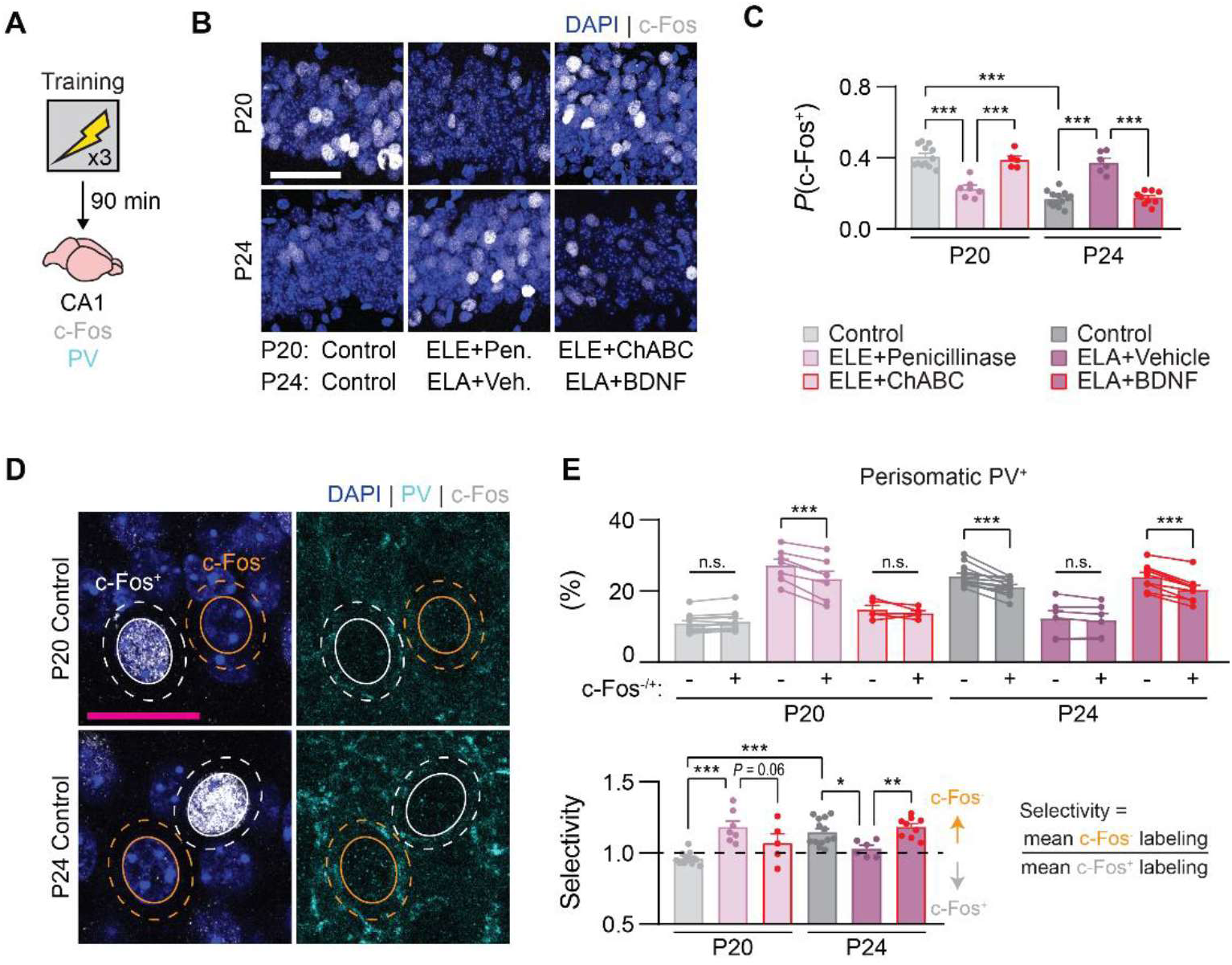
Early-life experience shapes trajectory for neuronal allocation to sparse engrams via PNNs. (A) Mice were trained in a contextual fear conditioning task on P20 or P24 and their brains were removed for c-Fos and PV immunohistochemistry. (B) Labeling of c-Fos^+^ nuclei (white) in the dorsal CA1 pyramidal layer. (C) Sparse expression of c-Fos in dorsal CA1 was observed in mice with mature PNNs (P20 ELE+Penicillinase, P24 Control, P24 ELA+BDNF) but not in mice with immature PNNs (P20 Control, P20 ELE+ChABC, and P24 ELA+Vehicle) (ANOVA, age × group interaction: *F*_2,46_ = 69.25, *P* < 0.00001). (D) Labeling of PV^+^ neurites (cyan) in the perisomatic region of putative engram (c-Fos^+^) and non-engram (c-Fos^-^) nuclei. (E) Following conditioning, PV^+^ neurites were more localized to the perisomatic region of non-engram cells in mice with mature PNNs (P20 ELE+Penicillinase, P24 Control, P24 ELA+BDNF) but not in mice with immature PNNs (P20 Control, P20 ELE+ChABC, and P24 ELA+Vehicle) (top, ANOVA, age × group × c-Fos interaction: *F*_2,46_ = 25.12, *P* < 0.00001; bottom, ANOVA, age × group interaction: *F*_2,46_ = 16.79, *P* < 0.00001). Data points represent individual mice with mean ± SEM. Scale bar, white: 50 µm, magenta: 20 µm. \**P* < 0.05, \*\**P* < 0.01, \*\*\**P* < 0.001, \*\*\*\**P* < 0.0001.

In typically developing mice, the accumulation of PNNs in dorsal CA1 increases the sparsity of engram ensembles representing an event by promoting structural and functional maturation of PV^+^ interneurons, which exclude neurons from engram ensembles via lateral inhibition following memory encoding (Morrison et al., 2016; Ramsaran et al., 2023; Rao-Ruiz et al., 2019; Rashid et al., 2016). We therefore examined the functional contribution of PV^+^ interneurons to the neuronal allocation process by measuring learning-induced localization of PV^+^ neurites to the perisomatic region of engram (c-Fos^+^) and non-engram (c-Fos^-^) cells in the dorsal CA1 pyramidal layer (**Figure 7D**). Consistent with the notion that sparse engram ensembles in dorsal CA1 require PV^+^ interneuron-mediated lateral inhibition, we observed greater localization of PV^+^ neurites in the perisomatic region of c-Fos^-^ non-engram cells (compared with c-Fos^+^ engram cells) in mice with high levels of PNNs (P20 ELE+Penicillinase, P24 Control, P24 ELA+BDNF; **Figure 7E**). These results demonstrate that early experiences, via their modulation of PNN growth, shape the trajectory for neuronal allocation to a sparse engram ensemble, which is required for precise memory formation.

## DISCUSSION

Here we performed a series of experiments to determine the role of postnatal experience in shaping the ability of the developing hippocampus to form precise episodic-like memories. By leveraging early-life experiential interventions known to affect brain maturation, we found that the onset of precise episodic-like memory formation in mice is influenced by early postnatal experience, rather than chronological age alone (**Figure 8**). Specifically, we found that ELA in the form of repeated bouts of maternal separation decelerated (but did not abolish) the maturation of PNNs in hippocampal subfield CA1 and the emergence of sparse engram ensembles and precise memory formation. By contrast, ELE, via physical and/or social enrichment, accelerated the maturation of CA1 PNNs and the emergence of sparse engram ensembles and precise memory formation. Moreover, the effects of ELA and ELE on the development of neural and behavioral memory precision could be countered by targeted treatments that reversed the post-intervention maturational state of the extracellular matrix in CA1. While ELA and ELE may have broad effects on neurodevelopmental trajectories for social, emotional, and cognitive development (Bale et al., 2010; Birnie & Baram, 2022; Opendak et al., 2017), this pattern of results suggests that they influence the developmental onset of precise episodic-like memory formation by specifically modulating the pace of PNN maturation in CA1 subfield of the hippocampus.

**Figure 8.**
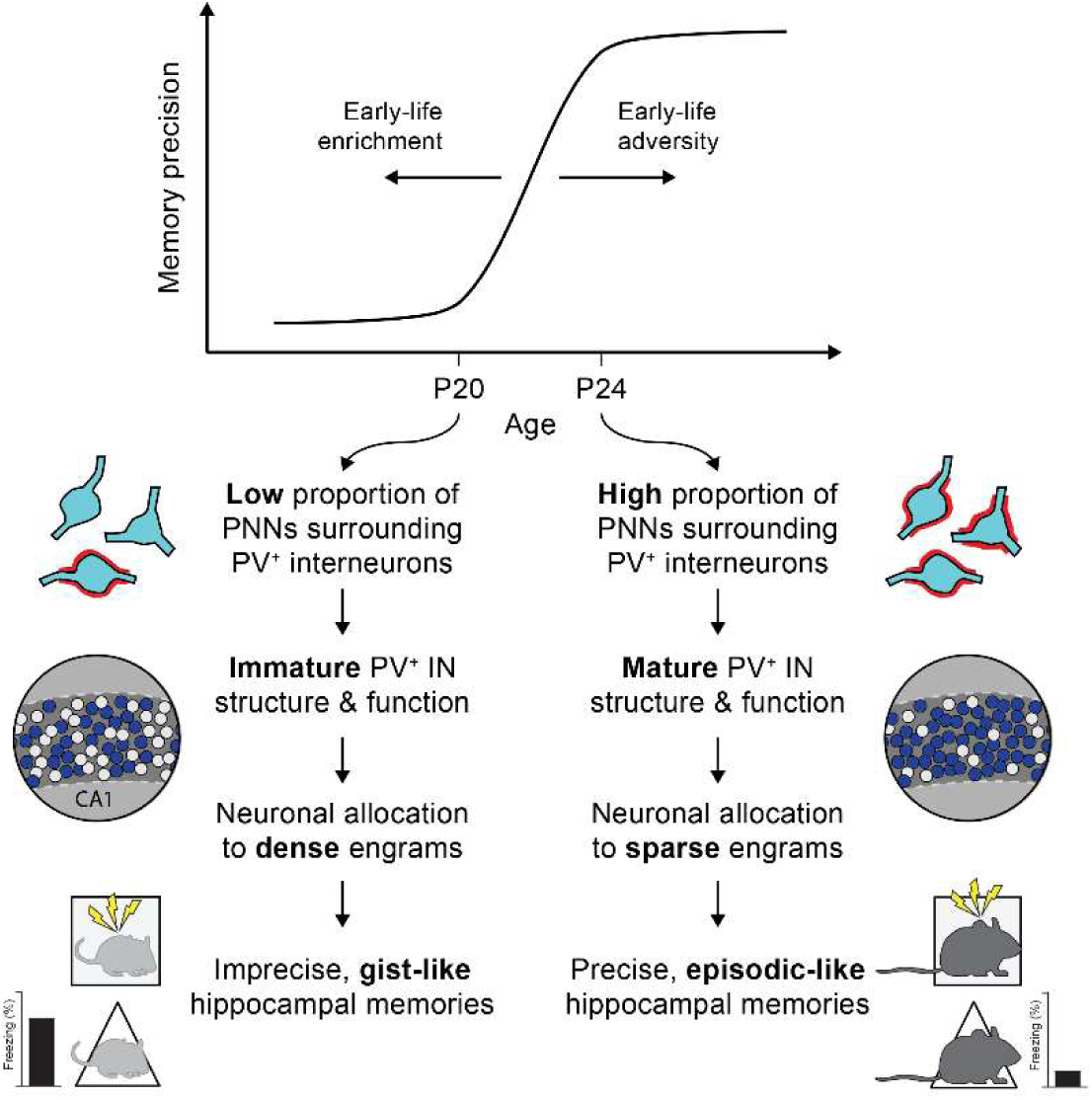
Early-life experiences determine the timing of episodic-like memory development in mice. The precision of event memories increases by P24, signaling the emergence of episodic-like memory function in mice. The development of episodic-like memory is supported by the maturation of extracellular PNNs around PV^+^ interneurons in CA1, which enable encoding of events by sparse neuronal engram ensembles. This developmental trajectory is decelerated in mice experiencing early-life adversity and is accelerated in mice experiencing early-life enrichment.

### A sensitive, but not critical, period for episodic-like memory development

Episodic-like memory is influenced by developmental processes occurring both pre- and postnatally. Embryonic neurogenesis configures the connectivity (Deguchi et al., 2011) and function (Huszár et al., 2022; Kveim et al., 2024) of neuronal ensembles in the mature hippocampus that support episodic-like memory. Yet although there is ample evidence to suggest that postnatal experiences impact episodic-like memory processes in adulthood (Hebb, 1947; Ivy et al., 2020; Molet et al., 2016; Woodcock & Richardson, 2000), it has remained unclear if and how sensitive the emergence of episodic-like memory during early life is to postnatal experiences. The emergence of several physiological correlates of hippocampal memory (e.g., place cells, theta sequences, sequential replay) studied in the dorsal CA1 of developing rats were previously shown to be unaffected by relevant spatial experiences, when those experiences occurred 0-1 days before the target electrophysiological recordings (Farooq & Dragoi, 2019; Wills et al., 2010). By contrast, recent studies using behavior to examine the development of persistent, hippocampus-dependent fear or object-location memories found that previous learning improves the persistence of a second related memory in juvenile rats and mice, if the related memory is formed within 24-48 h of the first experience (Bessières et al., 2020; Contreras et al., 2023). Therefore, previous hippocampal learning can influence later memory function in juvenile rodents, but only within a brief temporal window and when there are commonalities between the sequential experiences.

Previously, we found that rapid hippocampal learning (i.e., contextual fear conditioning) did not induce PNN formation in juvenile mice (Ramsaran et al., 2023). In the current study, we found that adverse or enriching experiences in the home environment influenced the precision of hippocampus-dependent contextual fear memories formed on P20 or P24, despite these experiences being not directly related to the fear memories. Therefore, whether postnatal experiences mature PNNs in the hippocampus likely depends on the type, timing, and/or duration of the experiences. Notably, experiences occurring during the first ∼4 postnatal weeks in mice are forgotten rapidly (over the course of hours and days). Offline processing of memories for experiences occurring during the ELA and ELE periods modulate the timing of hippocampal PNN maturation. Alternatively, ELA and ELE might modulate hippocampal PNN maturation by transiently altering neuronal activity within the maturing hippocampal-cortical circuitry, which is required for hippocampal circuit development (Donato et al., 2017) and PNN formation (Devienne et al., 2021).

Contrary to a recent proposal (Alberini & Travaglia, 2017; Travaglia et al., 2016), our findings support the notion that the early postnatal hippocampus in mice undergoes a sensitive period, but not critical period, for the maturation of episodic-like memory function. Critical periods are defined by the necessity of specific postnatal experiences in directing typical development, thereby establishing lifelong neural circuit performance. Accordingly, depriving animals of the requisite experiences (e.g., vision) during critical periods results in deficits in neural circuit performance (e.g., amblyopia) that persist into adulthood (Birch, 2013; Espinosa & Stryker, 2012; Holmes et al., 2011; Iny et al., 2006; Morishita & Hensch, 2008). In comparison, during sensitive periods, experience can exert powerful control over (but is not necessary for) normal neural circuit maturation. In our study, we found that deprivation by ELA delayed the formation of CA1 PNNs and the onset of memory precision, such that impairments in CA1 PNN growth and memory precision were observed in ELA mice on P24, but not P60. Therefore, in the weeks following cessation of ELA, hippocampal maturation **can “catch up” with that of control** mice. Consistent with this interpretation, anatomical maturation of the hippocampus can be rescued in developing mice with temporarily suppressed hippocampal-cortical activity by removing the suppression of neuronal activity (Donato et al., 2017), and functional maturation of hippocampal neuronal ensembles can be rescued in rats raised in sphere environments by repeatedly exposing them to linear environments (Farooq & Dragoi, 2023). Whether more severe forms of postnatal deprivation in mice would result in lifelong impairments in CA1 PNNs and memory precision indicative of a critical period, or whether the postnatal period is a critical (rather than sensitive) period for episodic-like memory in other species (Elliott & Richardson, 2019), remain open questions for future research. A more complete understanding of the hippocampal sensitive or critical period for episodic-like memory requires additional studies to define which parameters (e.g., types and amounts) of early experiences are most beneficial or deleterious for lifelong memory function. This may necessitate a granular approach, as hippocampal subfields with different developmental trajectories (e.g., CA1 vs. CA3 and DG; (Donato et al., 2017; Keresztes et al., 2018; Lavenex et al., 2006; Ramsaran et al., 2023)) may have different sensitivities to different types and time windows of postnatal experiences.

### Conserved mechanisms for experience-dependent neural circuit maturation across sensory and cognitive systems

In our previous study, we demonstrated that the emergence of episodic-like memory precision in mice requires the development of perisomatic inhibition in CA1, which relies on age-dependent formation of PNNs surrounding PV^+^ interneurons (Ramsaran et al., 2023). The same molecular and cellular mechanisms in V1 are responsible for an age-dependent increase in visual acuity, albeit at a later chronological age (Carulli et al., 2010; Espinosa & Stryker, 2012; Hensch, 2005; Huang et al., 1999; Pizzorusso et al., 2002). This suggests that PNN-dependent maturation of inhibitory circuits is a conserved mechanism for neural circuit maturation across sensory and cognitive systems in the brain.

In the current study, we further show that, similar to PNNs in V1, the timing of PNN formation in dorsal CA1 is controlled by early life experiences. Experience-dependent formation of PNNs in the hippocampus may shift the trajectory for episodic-like memory by improving neural representations of memory-related stimuli, similar to the sharpening of neuronal tuning to visual stimuli in V1 following the critical period (Huang et al., 1999; Tan et al., 2020). To our knowledge, one study has examined the effects of early-life experiences on the development of hippocampal neurons’ representation of space (Farooq & Dragoi, 2023). Here, the authors report that in the hippocampus of developing rats raised in spheres (i.e., environments that lack crucial features of Euclidean geometry), place cells are less precise and there is a reduced repertoire of neuronal ensembles available to encode experiences (versus control rats raised in cuboid environments), suggesting that this form of deprivation may also delay the maturation of episodic-like memory. Whether sphere-rearing affects PNN maturation in the hippocampus is unknown, but this is an intriguing possibility given that reduced precision of spatial representations is observed upon PNN digestion in the entorhinal cortex of adult rats (Christensen et al., 2021). Relatedly, in our previous work we showed that memory engrams, defined as neuronal ensembles encoding a given memory, become sparser in the dorsal CA1 with the maturation of PNNs and PV^+^ interneurons (Ramsaran et al., 2023). Extending these findings, we observed that the density of neuronal engram ensembles in dorsal CA1 of mice reared under different experiential conditions tracked the maturational state of local PNNs and PV+ interneurons, rather than chronological age. These data suggest that impairments in memory-guided behavior following alterations in postnatal experience may stem from deficiencies in neural coding, which are controlled by PNNs.

Is experience-dependent growth of PNNs a brain-wide mechanism for neural circuit maturation? In addition to their established roles in sensory cortices, in many brain regions including subdivisions of the medial prefrontal cortex (Baker et al., 2017), the amygdala (Gogolla et al., 2009), nucleus accumbens (Nardou et al., 2023), and hypothalamus (Mirzadeh et al., 2019), PNNs are implicated in neural circuit maturation. However, causal links between postnatal experience, PNN formation, and behavior have not been established in most cases. Importantly, the effects of enriching and adverse experiences on PNN formation and stability are age- and region-specific. For example, while ELE was shown to promote PNN formation in CA1 in the current study, the same intervention during adulthood led to reduced brevican-rich PNNs in CA1 (Favuzzi et al., 2017). Moreover, while ELA was shown to delay PNN formation in CA1 in the current study, different forms of ELA promote PNN formation in the amygdala (Gildawie et al., 2020; Guadagno et al., 2020) and anterior cingulate cortex (Catale et al., 2022) of rodents. The former of these is thought to contribute to the accelerated (rather than decelerated) emergence of adult-like extinction learning (Callaghan & Richardson, 2011), which requires PNNs in the amygdala (Gogolla et al., 2009). If PNN-dependent increases in inhibition regulate neural circuit and behavioral maturation across different regions of the brain, it is clear that the impact of specific postnatal experiences on PNN developmental trajectories for brain maturation are not uniform, as proposed recently (Tooley et al., 2021).

### Implications for episodic memory development in humans

Whether the human hippocampus undergoes a sensitive or critical period for maturation of episodic memory function remains an open question. Studies of individuals with varied childhood histories, including differences in family socioeconomic status (SES; which associates with the number and severity of stressful life events, (Evans & English, 2002)), suggest that, as in rodent species, adverse or enriching experiences during infancy and childhood can result in early differences in hippocampus-dependent memory which may persist into adulthood. For example, children that have experienced numerous, often severe, adverse life events (e.g., physical and/or sexual abuse, or adversity associated with institutionalized care or non-nurturing or -responsive parental care) and children from low-SES families develop smaller hippocampi and perform more poorly in episodic or spatial memory tasks compared to children without a history of adversity and children from high-SES families (Barch et al., 2019; Bremner, 2003; Decker et al., 2020; Hackman et al., 2010; Humphreys et al., 2019; Levine et al., 2005). Conversely, enriching childhood experiences (e.g., early childhood education, or enrichment associated with nurturing or responsive parental care) can promote hippocampal growth, improve memory, and mitigate cognitive deficits and other negative health outcomes following childhood adversity (F. Campbell et al., 2014; F. A. Campbell et al., 2001; Luby et al., 2012; Nelson et al., 2007).

The current study suggests that these experiences during the first years of life may alter the developmental trajectory of human hippocampal PNNs and PV^+^ interneurons (Lehner et al., 2024), providing a potential neurobiological mechanism by which different childhood experiences can shape lifelong episodic memory function. PNN maturation is dysregulated in cortex of individuals that experienced childhood adversity (Tanti et al., 2022), but whether similar dysregulation occurs in the human hippocampus is unknown. Understanding the effects early experiences have on the developing brain will lead to better-informed policy and practices aimed at preventing, minimizing, or mitigating the consequences of childhood adversity and bolstering the gains made with childhood enrichment.

## MATERIALS AND METHODS

### Mice

All procedures were approved by the Animal Care and Use Committee at the Hospital for Sick Children and were conducted in accordance with Canadian Council on Animal Care and National Institutes of Health guidelines. For all experiments, male and female wild-type mice derived from a C57BL/6N and 129S6 cross (Taconic Farms, Germantown, NY) were used. All mice were bred at the Hospital for Sick Children and maintained on a 12 h light/dark cycle (lights on at 0700 h), with ad libitum access to food and water. Litter birthdate was designated postnatal day (P) 0. Litter sizes ranged in size from 5-12 pups, and when possible, pups were cross-fostered to different lactating dams on P1-3 to equalize litter sizes and increase experiential diversity of offspring. Except where noted (see **Experiential Interventions**), experimental mice were housed in breeding cages (identical to standard housing cages) with the dam and male breeder from P0-20 and then weaned on P21 with same-sex littermates into new standard housing cages, with 2-5 mice per cage. Mice used for behavioral experiments on P20-21 remained in the breeding cage for the duration of the study. Each experimental condition contained mice derived from 2-6 separate litters.

### Experiential Interventions

Two experiential interventions were used, referred to hereafter as early-life adversity (ELA) and early-life enrichment (ELE). The ELA protocol was adapted from previous studies using C57BL/6J mice (George et al., 2010; Murthy et al., 2019). ELA consisted of maternal separation for 6 h per day from P6-16 (inclusive) followed by early weaning on P17. During separation, all pups from the litter were removed from the breeding cage and placed into a new cage with fresh bedding, and the cage of separated pups was moved to a different room from the breeding cage and placed on a heating pad turned to the lowest setting. From P12 onwards, the separated pups had free access to food and water, although there was little to no consumption of the solid food, which is usually not eaten before ∼P17 (König & Markl, 1987). On P17, ELA mice were weaned to new cages with same-sex littermates as described above. Control mice did not experience maternal separation and were weaned typically on P21.

The ELE protocol was adapted from previous studies using mice (Sztainberg & Chen, 2010). For the ELE intervention, all pups from the litter and the dam were relocated to a larger polycarbonate rat cage (enrichment cage) beginning on P6. The enrichment cages contained 3 huts and/or domes, 3-4 pieces of commercially purchased hamster tunnels, and ample nesting materials (twice the amount provided in standard cages).

Enrichment was rearranged or changed every ∼3 days, and enrichment cages were changed once per week. ELE mice were housed in the enrichment cages until P19 (or P18 for mice undergoing surgery, **Figure 6**), when they were returned to standard housing conditions before experiments began on P20. Control mice remained in standard housing cages during the equivalent period.

A subset of ELA, ELE, and control mice from were weighed on P6, P13, P20, P24, and P60 and observed for eye opening on P11-14. Offspring survival was not affected by the experiential interventions, and there were no observable changes in mouse health in any of the conditions, other than modest changes in weight gain in ELE and ELA mice (**Figure S1**).

### Surgery

Reagents used for intracranial injections were dissolved in solution using the appropriate vehicle, divided into 5 µl aliquots, and stored at -80 °C until use. Recombinant brain-derived neurotrophic factor (BDNF; Peprotech, cat# 450-02) was dissolved in phosphate-buffered saline (PBS) at a concentration of 0.33 µg/ml. Chondroitinase ABC (ChABC) from *Proteus vulgaris* (Sigma, cat# C3667) and penicillinase from *Bacillus cereus* (Sigma, cat# P0389) were separately dissolved in 0.1% bovine serum albumin in PBS at a concentration of 50 U/ml.

Surgeries occurred on P18 and P21 for experiments using P20 and P24 mice, respectively. Mice were pretreated with atropine sulfate (0.1 mg/kg, i.p.), anesthetized with isoflurane (3% induction, 1.0-1.5% maintenance), and administered meloxicam (4 mg/kg, s.c.) for analgesia. Mice were placed into stereotaxic frames and topically administered lidocaine around the incision site. The scalp was incised and retracted before drilling two holes above the dorsal CA1 (AP -1.7 mm and ML ±1.35 mm from bregma). A glass micropipette connected to a Hamilton microsyringe was filled with the appropriate solution and then lowered to the CA1 pyramidal cell layer (DV -1.45 mm from bregma). Solutions were injected at a rate of 0.1 µl/min to a total volume of 0.75 µl. The glass micropipette was left in place for an additional 10 min after the microinjection. Following both microinjections, the incision was closed using sutures, polysporin was applied to the wound, and mice were administered 0.5 ml (s.c.) 0.9% saline. Recovering mice were placed in a clean cage on a heating pad until ambulatory, at which point they were returned to the colony. Animals were monitored twice daily until the end of the experiment.

### Behavior

#### Contextual fear conditioning

Fear conditioning was performed as previously described (Ramsaran et al., 2023). At the appropriate postnatal age (P20 or P24), mice were placed in 31 × 24 × 21 cm metallic test chambers equipped with shock-grid floors (Med Associates). On the training day, mice remained in the chamber for a total of 5 min, and three 0.5 mA foot shocks (0.5 mA, 2 s duration) were delivered at 120, 180, and 240 s. The following day, mice were placed back into the same chamber (Context A) or a similar but novel chamber (Context B) for the contextual fear memory test (5 min), during which no additional stimuli were presented. Context B was similar in size to Context A, but differed in shape, color, texture, and location. Context B chambers were in a room adjacent to the Context A chambers, and smooth white plastic inserts were used to create triangular walls and to cover the shock-grid floor. Mouse behavior during the training and test sessions was recorded using overhead cameras and FreezeFrame v.3.32 software (Actimetrics). We used freezing behavior in each context (defined as the cessation of all movement, except for breathing (Fanselow, 1980)) as a proxy for contextual fear memory recall. The amount of time mice spent freezing during each session was scored semi-automatically using FreezeFrame.

#### Open field

We examined anxiety-like phenotype of adult mice using an open field test, as previously described (Ramsaran et al., 2023). Mice were placed in the center of a 45 × 45 × 20 cm open field arena located in a dimly lit room and allowed to freely explore for 10 min. Mouse locomotion was tracked using overhead cameras and Limelight software (Actimetrics). The total distance traveled and amount of time spent in the center of the arena was scored automatically using Limelight.

### Histology

Procedures were conducted as described previously (Ramsaran et al., 2023). At the appropriate age, mice were transcardially perfused with chilled PBS followed by 4% paraformaldehyde (PFA). Volumes were adjusted based on mouse age. Brains were removed to conical tubes containing PFA and post-fixed overnight at 4 °C, followed by 30% sucrose solution until submerged (48-72 h). Brains were sectioned coronally into 50-µm sections across the anterior-posterior extent of the dorsal hippocampus using a Lecia CM1850 and stored in PBS containing 0.1% NaN_3_ at 4 °C until staining.

Six to eight tissue sections (200-μm intervals) from each brain were used for immunohistochemistry. All incubations occurred at room temperature unless otherwise stated. Free-floating sections were blocked with PBS containing 4% normal goat serum and 0.5% Triton-X for 1 h, followed by incubation with biotin-conjugated lectin from *Wisteria floribunda* (WFA; 1:1000, Sigma, cat# L1516), rabbit anti-parvalbumin (1:1000, Swant, cat# PV27) or guinea-pig anti-parvalbumin (1:1000, Swant, cat# GP72), and guinea-pig anti-c-Fos (1:1000, Synaptic Systems, Cat# 226-308) primary antibodies in fresh blocking solution overnight at 4 °C. Sections were washed three times for 15 min each in PBS containing 0.1% Tween-20 (PBST) and then incubated with secondary antibodies streptavidin Alexa Fluor 568 (1:500, Invitrogen, cat# S-11226) or streptavidin Alexa Fluor 647(1:500, Invitrogen, cat# S-32357), goat anti-rabbit Alexa Fluor 488 (1:500, Invitrogen, cat# A-11008) or goat anti-guinea-pig Alexa Fluor 488 (1:500, Invitrogen, cat# A-11073), and goat anti-guinea-pig Alexa Fluor 647 (for c-Fos; 1:500, Initrogen,cat# A-21450) in PBST for 2 h. Sections were washed with PBS, counterstained with DAPI (1:10000), mounted on gel-coated slides, and coverslipped with Permafluor mounting medium (ThermoFisher Scientific, cat# TA-030-FM).

Images were taken on a confocal laser scanning microscope (LSM 710; Zeiss) using a 40× objective with numerical aperture of 1.3 (for PNN and PV^+^ soma quantification) or 63× objective with numerical aperture of 1.4 (for c-Fos and perisomatic PV^+^ quantification). For image quantification, 4-6 z-stacks per brain were obtained (15-30 slices, 1-µm step size). All images for a given experiment were obtained in a single session using identical laser power, photomultiplier gain, pinhole, and detection filter settings.

Quantification of WFA^+^, PV^+^, and c-Fos^+^ cells within the acquired z-stacks was performed manually using Fiji (National Institutes of Health). Cells per tissue volume (mm^-3^) was calculated for each marker and values corresponding to images obtained from the same mouse were averaged. Quantification of PV^+^ neurites in the perisomatic regions of c-Fos^-^ and c-Fos^+^ cells in the CA1 pyramidal layer was performed using Fiji as previously described (Davis et al., 2017; Ramsaran et al., 2023; Trouche et al., 2013). Briefly, PV^+^ immunofluorescence was binarized using a gray value threshold that was manually determined from images from a P24 Control mouse, and the same threshold was applied to all images regardless of mouse group. Next, 50-80 nuclei expressing c-Fos (c-Fos^+^) or not (c-Fos^-^) were outlined in Fiji and the percentage of area covered by (binarized) PV^+^ labeling within a 3-μm band was measured. Perisomatic PV^+^ labeling was averaged for each mouse for c-Fos^+^ and c-Fos^-^ nuclei separately. Selectivity of PV+ labeling was calculated for each mouse as (mean c-Fos^-^ labeling / mean c-Fos^+^ labeling).

### Statistical Analyses

No statistical tests were used to predetermine sample sizes, but sample sizes were similar to those reported previously (Akers et al., 2014; Ramsaran et al., 2023; Vetere et al., 2021). To achieve approximately equivalent group sizes, litters were pseudo-randomly assigned to the experiential interventions (ELA vs. Control or ELE vs. Control), and for surgical experiments, mice within each litter were pseudo-randomly assigned to different treatment conditions (Vehicle vs. BDNF or Penicillinase vs. ChABC). Experimenters were blinded to group assignments during data collection and quantification, except for testing context for the fear conditioning experiments. Statistical analyses were performed using Statistica software (Dell Inc. 2016, version 13) and Graphpad Prism (version 8.0.1). Data were analyzed unpaired t-tests or analysis of variance (ANOVA), followed by Newman-Keuls post-hoc comparisons. Statistical significance was set at P < 0.05. We did not include sex as a factor in the analyses (other than weight; **Figure S1**), but there were no apparent sex differences on the key neural and behavioral measures (**Figure S2**). For the weight analyses, we did not include sex as a factor for P6, P13, or P24 time points, as there are no weight differences between male and female prepubescent mice.

## Author Contributions

A.I.R. and P.W.F. designed the experiments. A.I.R., S.V., J.G., and M.L.D. conducted the experiments. A.I.R. analyzed the data and wrote the first draft of the paper. All authors edited the paper.

## Acknowledgements

We thank A. DeCristofaro, D. Lin, and M. Yamamato for laboratory support. This work was supported by grants from the Canadian Institutes of Health Research (PJT180530 to P.W.F.), Brain Canada (to P.W.F. and S.A.J.) and the National Institutes of Health (R01 MH119421 to P.W.F. and S.A.J.). A.I.R. was supported by the National Institutes of Health (F31 MH120920) and the Natural Sciences and Engineering Research Council of Canada. M.L.D. was supported by a CIHR Vanier Canada graduate scholarship (M.L.D.).

---

**Figure S1. Effects of early-life experiences on weight gain and eye opening.**

(A) Early-life adversity and enrichment produce modest changes in body weight compared to control conditions (P6 to P20, RM-ANOVA, age × experience interaction: *F*_4,126_ = 26.99, *P* < 0.0001; P24, unpaired *t*-test, *t*_41_ = 5.31, *P* < 0.0001; P60-F, unpaired *t*-test, *t*_13_ = 0.30, *P* = 0.77; P60-M, unpaired *t*-test, *t*_16_ = 2.66, *P* < 0.05). (B) Early-life adversity and enrichment do not alter the age of eye opening (ANOVA, no main effect of experience: *F*_2,62_ = 2.19, *P* = 0.12). Data points represent individual mice with mean ± SEM. \**P* < 0.05, \*\**P* < 0.01, \*\*\**P* < 0.001, \*\*\*\**P* < 0.0001.

**Figure S2. No sex differences in the effects of early-life experiences on trajectories for PNN formation and memory precision.**

(A) The effects of early-life adversity on levels of CA1 WFA^+^ PNNs and PNN-enwrapped PV^+^ interneurons in P24 mice was not dependent on sex. (B) Experimental schedule for assessing memory precision. Mice were trained in a contextual fear conditioning task on P24, and 1 d later, they were tested in the same context (Context A) or a similar, novel context (Context B). (C) The effects of early-life adversity on memory precision in P24 mice were not dependent on sex. (D) The effects of early-life enrichment on levels of CA1 WFA^+^ PNNs and PNN-enwrapped PV^+^ interneurons in P20 mice was not dependent on sex. (E) Experimental schedule for assessing memory precision. Mice were trained in a contextual fear conditioning task on P20, and 1 d later, they were tested in the same context (Context A) or a similar, novel context (Context B). (F) The effects of early-life enrichment on memory precision in P20 mice were not dependent on sex. Data points represent individual mice with mean ± SEM. Data shown in panels 2A,C and 2D,F are reproduced from Figures 2C,G and 5C,G, respectively, and stratified by sex. \**P* < 0.05, \*\**P* < 0.01, \*\*\**P* < 0.001, \*\*\*\**P* < 0.0001.

## References

Akers, K. G., Martinez-Canabal, A., Restivo, L., Yiu, A. P., De Cristofaro, A., Hsiang, H.-L. (Liz), Wheeler, A. L., Guskjolen, A., Niibori, Y., Shoji, H., Ohira, K., Richards, B. A., Miyakawa, T., Josselyn, S. A., & Frankland, P. W. (2014). Hippocampal Neurogenesis Regulates Forgetting During Adulthood and Infancy. Science, 344(6184), 598–602. 10.1126/science.1248903

Alberini, C. M., & Travaglia, A. (2017). Infantile Amnesia: A Critical Period of Learning to Learn and Remember. Journal of Neuroscience, 37(24), 5783–5795. 10.1523/JNEUROSCI.0324-17.2017

Amso, D., Salhi, C., & Badre, D. (2019). The relationship between cognitive enrichment and cognitive control: A systematic investigation of environmental influences on development through socioeconomic status. Developmental Psychobiology, 61(2), 159–178. 10.1002/dev.21794

Anderson, M. J., & Riccio, D. C. (2005). Ontogenetic forgetting of stimulus attributes. Learning & Behavior, 33(4), 444–453. 10.3758/BF03193183

Baker, K. D., Gray, A. R., & Richardson, R. (2017). The development of perineuronal nets around parvalbumin gabaergic neurons in the medial prefrontal cortex and basolateral amygdala of rats. Behavioral Neuroscience, 131(4), 289–303. 10.1037/bne0000203

Bale, T. L., Baram, T. Z., Brown, A. S., Goldstein, J. M., Insel, T. R., McCarthy, M. M., Nemeroff, C. B., Reyes, T. M., Simerly, R. B., Susser, E. S., & Nestler, E. J. (2010). Early Life Programming and Neurodevelopmental Disorders. Biological Psychiatry, 68(4), 314–319. 10.1016/j.biopsych.2010.05.028

Barch, D. M., Harms, M. P., Tillman, R., Hawkey, E., & Luby, J. L. (2019). Early childhood depression, emotion regulation, episodic memory, and hippocampal development. Journal of Abnormal Psychology, 128(1), 81–95. 10.1037/abn0000392

Bessières, B., Travaglia, A., Mowery, T. M., Zhang, X., & Alberini, C. M. (2020). Early life experiences selectively mature learning and memory abilities. Nature Communications, 11(1), Article 1. 10.1038/s41467-020-14461-3

Birch, E. E. (2013). Amblyopia and binocular vision. Progress in Retinal and Eye Research, 33, 67–84. 10.1016/j.preteyeres.2012.11.001

Birnie, M. T., & Baram, T. Z. (2022). Principles of emotional brain circuit maturation. Science, 376(6597), 1055–1056. 10.1126/science.abn4016

Bremner, J. D. (2003). Long-term effects of childhood abuse on brain and neurobiology. Child and Adolescent Psychiatric Clinics, 12(2), 271–292. 10.1016/S1056-4993(02)00098-6

Callaghan, B. L., & Richardson, R. (2011). Maternal separation results in early emergence of adult-like fear and extinction learning in infant rats. Behavioral Neuroscience, 125(1), 20–28. 10.1037/a0022008

Campbell, F. A., Pungello, E. P., Miller-Johnson, S., Burchinal, M., & Ramey, C. T. (2001). The development of cognitive and academic abilities: Growth curves from an early childhood educational experiment. Developmental Psychology, 37(2), 231–242. 10.1037/0012-1649.37.2.231

Campbell, F., Conti, G., Heckman, J. J., Moon, S. H., Pinto, R., Pungello, E., & Pan, Y. (2014). Early Childhood Investments Substantially Boost Adult Health. Science, 343(6178), 1478–1485. 10.1126/science.1248429

Cancedda, L., Putignano, E., Sale, A., Viegi, A., Berardi, N., & Maffei, L. (2004). Acceleration of Visual System Development by Environmental Enrichment. Journal of Neuroscience, 24(20), 4840–4848. 10.1523/JNEUROSCI.0845-04.2004

Carulli, D., Pizzorusso, T., Kwok, J. C. F., Putignano, E., Poli, A., Forostyak, S., Andrews, M. R., Deepa, S. S., Glant, T. T., & Fawcett, J. W. (2010). Animals lacking link protein have attenuated perineuronal nets and persistent plasticity. Brain, 133(8), 2331–2347. 10.1093/brain/awq145

Carulli, D., & Verhaagen, J. (2021). An Extracellular Perspective on CNS Maturation: Perineuronal Nets and the Control of Plasticity. International Journal of Molecular Sciences, 22(5), Article 5. 10.3390/ijms22052434

Catale, C., Martini, A., Piscitelli, R. M., Senzasono, B., Iacono, L. L., Mercuri, N. B., Guatteo, E., & Carola, V. (2022). Early-life social stress induces permanent alterations in plasticity and perineuronal nets in the mouse anterior cingulate cortex. European Journal of Neuroscience, 56(10), 5763–5783. 10.1111/ejn.15825

Chapman, B., & Stryker, M. P. (1993). Development of orientation selectivity in ferret visual cortex and effects of deprivation. Journal of Neuroscience, 13(12), 5251 5262. 10.1523/JNEUROSCI.13-12-05251.1993

Christensen, A. C., Lensjø, K. K., Lepperød, M. E., Dragly, S.-A., Sutterud, H., Blackstad, J. S., Fyhn, M., & Hafting, T. (2021). Perineuronal nets stabilize the grid cell network. Nature Communications, 12(1), Article 1. 10.1038/s41467-020-20241-w

Contreras, M. P., Fechner, J., Born, J., & Inostroza, M. (2023). Accelerating Maturation of Spatial Memory Systems by Experience: Evidence from Sleep Oscillation Signatures of Memory Processing. Journal of Neuroscience, 43(19), 3509–3519. 10.1523/JNEUROSCI.1967-22.2023

Davis, P., Zaki, Y., Maguire, J., & Reijmers, L. G. (2017). Cellular and oscillatory substrates of fear extinction learning. Nature Neuroscience, 20(11), Article 11. 10.1038/nn.4651

Daw, N. W., Fox, K., Sato, H., & Czepita, D. (1992). Critical period for monocular deprivation in the cat visual cortex. Journal of Neurophysiology, 67(1), 197–202. 10.1152/jn.1992.67.1.197

Decker, A. L., Duncan, K., Finn, A. S., & Mabbott, D. J. (2020). Children’s family income is associated with cognitive function and volume of anterior not posterior hippocampus. Nature Communications, 11(1), Article 1. 10.1038/s41467-020-17854-6

Deguchi, Y., Donato, F., Galimberti, I., Cabuy, E., & Caroni, P. (2011). Temporally matched subpopulations of selectively interconnected principal neurons in the hippocampus. Nature Neuroscience, 14(4), Article 4. 10.1038/nn.2768

Devienne, G., Picaud, S., Cohen, I., Piquet, J., Tricoire, L., Testa, D., Nardo, A. A. D., Rossier, J., Cauli, B., & Lambolez, B. (2021). Regulation of Perineuronal Nets in the Adult Cortex by the Activity of the Cortical Network. Journal of Neuroscience, 41(27), 5779–5790. 10.1523/JNEUROSCI.0434-21.2021

Donato, F., Jacobsen, R. I., Moser, M.-B., & Moser, E. I. (2017). Stellate cells drive maturation of the entorhinal-hippocampal circuit. Science, 355(6330), eaai8178. 10.1126/science.aai8178

Duffy, S. N., Craddock, K. J., Abel, T., & Nguyen, P. V. (2001). Environmental Enrichment Modifies the PKA-Dependence of Hippocampal LTP and Improves Hippocampus-Dependent Memory. Learning & Memory, 8(1), 26–34. 10.1101/lm.36301

Elliott, N. D., & Richardson, R. (2019). The effects of early life stress on context fear generalization in adult rats. Behavioral Neuroscience, 133(1), 50–58. 10.1037/bne0000289

Erzurumlu, R. S., & Gaspar, P. (2012). Development and critical period plasticity of the barrel cortex. European Journal of Neuroscience, 35(10), 1540–1553. 10.1111/j.1460-9568.2012.08075.x

Espinosa, J. S., & Stryker, M. P. (2012). Development and Plasticity of the Primary Visual Cortex. Neuron, 75(2), 230–249. 10.1016/j.neuron.2012.06.009

Evans, G. W., & English, K. (2002). The Environment of Poverty: Multiple Stressor Exposure, Psychophysiological Stress, and Socioemotional Adjustment. Child Development, 73(4), 1238–1248. 10.1111/1467-8624.00469

Evans, G. W., & Schamberg, M. A. (2009). Childhood poverty, chronic stress, and adult working memory. Proceedings of the National Academy of Sciences, 106(16), 6545–6549. 10.1073/pnas.0811910106

Fanselow, M. S. (1980). Conditional and unconditional components of post-shock freezing. The Pavlovian Journal of Biological Science: Official Journal of the Pavlovian, 15(4), 177–182. 10.1007/BF03001163

Farooq, U., & Dragoi, G. (2019). Emergence of preconfigured and plastic time-compressed sequences in early postnatal development. Science, 363(6423), 168–173. 10.1126/science.aav0502

Farooq, U., & Dragoi, G. (2023). Geometric experience sculpts the development and dynamics of hippocampal sequential cell assemblies (p. 2023.12.04.570026). bioRxiv. 10.1101/2023.12.04.570026

Favuzzi, E., Marques-Smith, A., Deogracias, R., Winterflood, C. M., Sánchez-Aguilera, A., Mantoan, L., Maeso, P., Fernandes, C., Ewers, H., & Rico, B. (2017). Activity-Dependent Gating of Parvalbumin Interneuron Function by the Perineuronal Net Protein Brevican. Neuron, 95(3), 639–655.e10. 10.1016/j.neuron.2017.06.028

Fawcett, J. W., Oohashi, T., & Pizzorusso, T. (2019). The roles of perineuronal nets and the perinodal extracellular matrix in neuronal function. Nature Reviews Neuroscience, 20(8), Article 8. 10.1038/s41583-019-0196-3

Fox, K. (1992). A critical period for experience-dependent synaptic plasticity in rat barrel cortex. Journal of Neuroscience, 12(5), 1826–1838. 10.1523/JNEUROSCI.12-05-01826.1992

Freedman, D. G., King, J. A., & Elliot, O. (1961). Critical period in the social development of dogs. Science, 133, 1016–1017. 10.1126/science.133.3457.1016

Frick, K. M., Stearns, N. A., Pan, J.-Y., & Berger-Sweeney, J. (2003). Effects of Environmental Enrichment on Spatial Memory and Neurochemistry in Middle-Aged Mice. Learning & Memory, 10(3), 187–198. 10.1101/lm.50703

George, E. D., Bordner, K. A., Elwafi, H. M., & Simen, A. A. (2010). Maternal separation with early weaning: A novel mouse model of early life neglect. BMC Neuroscience, 11(1), 123. 10.1186/1471-2202-11-123

Gianfranceschi, L., Siciliano, R., Walls, J., Morales, B., Kirkwood, A., Huang, Z. J., Tonegawa, S., & Maffei, L. (2003). Visual cortex is rescued from the effects of dark rearing by overexpression of BDNF. Proceedings of the National Academy of Sciences, 100(21), 12486–12491. 10.1073/pnas.1934836100

Gildawie, K. R., Honeycutt, J. A., & Brenhouse, H. C. (2020). Region-specific Effects of Maternal Separation on Perineuronal Net and Parvalbumin-expressing Interneuron Formation in Male and Female Rats. Neuroscience, 428, 23–37. 10.1016/j.neuroscience.2019.12.010

Gogolla, N., Caroni, P., Lüthi, A., & Herry, C. (2009). Perineuronal Nets Protect Fear Memories from Erasure. Science, 325(5945), 1258–1261. 10.1126/science.1174146

Goodwill, H. L., Manzano-Nieves, G., Gallo, M., Lee, H.-I., Oyerinde, E., Serre, T., & Bath, K. G. (2019). Early life stress leads to sex differences in development of depressive-like outcomes in a mouse model. Neuropsychopharmacology, 44(4), Article 4. 10.1038/s41386-018-0195-5

Gordon, J. A., & Stryker, M. P. (1996). Experience-Dependent Plasticity of Binocular Responses in the Primary Visual Cortex of the Mouse. Journal of Neuroscience, 16(10), 3274–3286. 10.1523/JNEUROSCI.16-10-03274.1996

Guadagno, A., Verlezza, S., Long, H., Wong, T. P., & Walker, C.-D. (2020). It Is All in the Right Amygdala: Increased Synaptic Plasticity and Perineuronal Nets in Male, But Not Female, Juvenile Rat Pups after Exposure to Early-Life Stress. Journal of Neuroscience, 40(43), 8276–8291. 10.1523/JNEUROSCI.1029-20.2020

Guayasamin, M., Depaauw-Holt, L. R., Adedipe, I. I., Ghenissa, O., Vaugeois, J., Duquenne, M., Rogers, B., Latraverse-Arquilla, J., Peyrard, S., Bosson, A., & Murphy-Royal, C. (2024). Early-life stress induces persistent astrocyte dysfunction resulting in fear generalisation. eLife, 13. 10.7554/eLife.99988.1

Hackman, D. A., Farah, M. J., & Meaney, M. J. (2010). Socioeconomic status and the brain: Mechanistic insights from human and animal research. Nature Reviews Neuroscience, 11(9), Article 9. 10.1038/nrn2897

Hanover, J. L., Huang, Z. J., Tonegawa, S., & Stryker, M. P. (1999). Brain-Derived Neurotrophic Factor Overexpression Induces Precocious Critical Period in Mouse Visual Cortex. Journal of Neuroscience, 19(22), RC40–RC40. 10.1523/JNEUROSCI.19-22-j0003.1999

Harrison, R. V., Gordon, K. A., & Mount, R. J. (2005). Is there a critical period for cochlear implantation in congenitally deaf children? Analyses of hearing and speech perception performance after implantation. Developmental Psychobiology, 46(3), 252–261. 10.1002/dev.20052

Hebb, D. O. (1947). The effects of early experience on problem-solving at maturity. American Psychologist, 2, 306–307.

Hensch, T. K. (2005). Critical period plasticity in local cortical circuits. Nature Reviews Neuroscience, 6(11), Article 11. 10.1038/nrn1787

Holmes, J. M., Lazar, E. L., Melia, B. M., Astle, W. F., Dagi, L. R., Donahue, S. P., Frazier, M. G., Hertle, R. W., Repka, M. X., Quinn, G. E., Weise, K. K., & Pediatric Eye Disease Investigator Group, for the. (2011). Effect of Age on Response to Amblyopia Treatment in Children. Archives of Ophthalmology, 129(11), 1451–1457. 10.1001/archophthalmol.2011.179

Huang, Z. J., Kirkwood, A., Pizzorusso, T., Porciatti, V., Morales, B., Bear, M. F., Maffei, L., & Tonegawa, S. (1999). BDNF Regulates the Maturation of Inhibition and the Critical Period of Plasticity in Mouse Visual Cortex. Cell, 98(6), 739–755. 10.1016/S0092-8674(00)81509-3

Humphreys, K. L., King, L. S., Sacchet, M. D., Camacho, M. C., Colich, N. L., Ordaz, S. J., Ho, T. C., & Gotlib, I. H. (2019). Evidence for a sensitive period in the effects of early life stress on hippocampal volume. Developmental Science, 22(3), e12775. 10.1111/desc.12775

Huot, R. L., Plotsky, P. M., Lenox, R. H., & McNamara, R. K. (2002). Neonatal maternal separation reduces hippocampal mossy fiber density in adult Long Evans rats. Brain Research, 950(1), 52–63. 10.1016/S0006-8993(02)02985-2

Huszár, R., Zhang, Y., Blockus, H., & Buzsáki, G. (2022). Preconfigured dynamics in the hippocampus are guided by embryonic birthdate and rate of neurogenesis. Nature Neuroscience, 25(9), Article 9. 10.1038/s41593-022-01138-x

Iny, K., Heynen, A. J., Sklar, E., & Bear, M. F. (2006). Bidirectional Modifications of Visual Acuity Induced by Monocular Deprivation in Juvenile and Adult Rats. Journal of Neuroscience, 26(28), 7368–7374. 10.1523/JNEUROSCI.0124-06.2006

Ivy, A. S., Yu, T., Kramár, E., Parievsky, S., Sohn, F., & Vu, T. (2020). A Unique Mouse Model of Early Life Exercise Enables Hippocampal Memory and Synaptic Plasticity. Scientific Reports, 10(1), Article 1. 10.1038/s41598-020-66116-4

Keresztes, A., Ngo, C. T., Lindenberger, U., Werkle-Bergner, M., & Newcombe, N. S. (2018). Hippocampal Maturation Drives Memory from Generalization to Specificity. Trends in Cognitive Sciences, 22(8), 676–686. 10.1016/j.tics.2018.05.004

König, B., & Markl, H. (1987). Maternal care in house mice: I. The weaning strategy as a means for parental manipulation of offspring quality. Behavioral Ecology and Sociobiology, 20(1), 1–9. 10.1007/BF00292161

Kveim, V. A., Salm, L., Ulmer, T., Lahr, M., Kandler, S., Imhof, F., & Donato, F. (2024). Divergent recruitment of developmentally defined neuronal ensembles supports memory dynamics. Science, 385(6710), eadk0997. 10.1126/science.adk0997

Lambert, H. K., Sheridan, M. A., Sambrook, K. A., Rosen, M. L., Askren, M. K., & McLaughlin, K. A. (2017). Hippocampal Contribution to Context Encoding across Development Is Disrupted following Early-Life Adversity. Journal of Neuroscience, 37(7), 1925–1934. 10.1523/JNEUROSCI.2618-16.2017

Lavenex, P., Banta Lavenex, P., & Amaral, D. G. (2006). Postnatal Development of the Primate Hippocampal Formation. Developmental Neuroscience, 29(1–2), 179–192. 10.1159/000096222

Lehner, A., Hoffmann, L., Rampp, S., Coras, R., Paulsen, F., Frischknecht, R., Hamer, H., Walther, K., Brandner, S., Hofer, W., Pieper, T., Reisch, L.-M., Bien, C. G., & Blumcke, I. (2024). Age-dependent increase of perineuronal nets in the human hippocampus and precocious aging in epilepsy. Epilepsia Open, 9(4), 1372 1381. 10.1002/epi4.12963

Lesuis, S.L., Brosens, N., Immerzeel, N., van der Loo, R. J., Mitrić, M., Bielefeld, P., Fitzsimons, C. P., Lucassen, P. J., Kushner, S. A., van den Oever, M. C., & Krugers, H. J. (2021). Glucocorticoids Promote Fear Generalization by Increasing the Size of a Dentate Gyrus Engram Cell Population. Biological Psychiatry, 90(7), 494–504. 10.1016/j.biopsych.2021.04.010

Levine, S. C., Vasilyeva, M., Lourenco, S. F., Newcombe, N. S., & Huttenlocher, J. (2005). Socioeconomic Status Modifies the Sex Difference in Spatial Skill. Psychological Science, 16(11), 841–845. 10.1111/j.1467-9280.2005.01623.x

Li, R., Wang, X., Lin, F., Song, T., Zhu, X., & Lei, H. (2020). Mapping accumulative whole-brain activities during environmental enrichment with manganese-enhanced magnetic resonance imaging. NeuroImage, 210, 116588. 10.1016/j.neuroimage.2020.116588

Lorenz, K. (1935). Der Kumpan in der Umwelt des Vogels. Der Artgenosse als auslösendes Moment sozialer Verhaltungsweisen. [The companion in the bird’s world. The fellow-member of the species as releasing factor of social behavior.]. Journal Für Ornithologie. Beiblatt. (Leipzig), 83, 137–213. 10.1007/BF01905355

Luby, J. L., Barch, D. M., Belden, A., Gaffrey, M. S., Tillman, R., Babb, C., Nishino, T., Suzuki, H., & Botteron, K. N. (2012). Maternal support in early childhood predicts larger hippocampal volumes at school age. Proceedings of the National Academy of Sciences, 109(8), 2854–2859. 10.1073/pnas.1118003109

Madsen, H. B., & Kim, J. H. (2016). Ontogeny of memory: An update on 40 years of work on infantile amnesia. Behavioural Brain Research, 298, 4–14. 10.1016/j.bbr.2015.07.030

Malave, L., van Dijk, M. T., & Anacker, C. (2022). Early life adversity shapes neural circuit function during sensitive postnatal developmental periods. Translational Psychiatry, 12(1), Article 1. 10.1038/s41398-022-02092-9

McRae, P. A., Rocco, M. M., Kelly, G., Brumberg, J. C., & Matthews, R. T. (2007). Sensory Deprivation Alters Aggrecan and Perineuronal Net Expression in the Mouse Barrel Cortex. Journal of Neuroscience, 27(20), 5405–5413. 10.1523/JNEUROSCI.5425-06.2007

Miranda, M., Kent, B. A., Morici, J. F., Gallo, F., Saksida, L. M., Bussey, T. J., Weisstaub, N., & Bekinschtein, P. (2018). NMDA receptors and BDNF are necessary for discrimination of overlapping spatial and non-spatial memories in perirhinal cortex and hippocampus. Neurobiology of Learning and Memory, 155, 337–343. 10.1016/j.nlm.2018.08.019

Mirzadeh, Z., Alonge, K. M., Cabrales, E., Herranz-Pérez, V., Scarlett, J. M., Brown, J. M., Hassouna, R., Matsen, M. E., Nguyen, H. T., Garcia-Verdugo, J. M., Zeltser, L. M., & Schwartz, M. W. (2019). Perineuronal net formation during the critical period for neuronal maturation in the hypothalamic arcuate nucleus. Nature Metabolism, 1(2), Article 2. 10.1038/s42255-018-0029-0

Molet, J., Maras, P. M., Kinney-Lang, E., Harris, N. G., Rashid, F., Ivy, A. S., Solodkin, A., Obenaus, A., & Baram, T. Z. (2016). MRI uncovers disrupted hippocampal microstructure that underlies memory impairments after early-life adversity. Hippocampus, 26(12), 1618–1632. 10.1002/hipo.22661

Morishita, H., & Hensch, T. K. (2008). Critical period revisited: Impact on vision. Current Opinion in Neurobiology, 18(1), 101–107. 10.1016/j.conb.2008.05.009

Morrison, D. J., Rashid, A. J., Yiu, A. P., Yan, C., Frankland, P. W., & Josselyn, S. A. (2016). Parvalbumin interneurons constrain the size of the lateral amygdala engram. Neurobiology of Learning and Memory, 135, 91–99. 10.1016/j.nlm.2016.07.007

Mowery, T. M., Kotak, V. C., & Sanes, D. H. (2015). Transient Hearing Loss Within a Critical Period Causes Persistent Changes to Cellular Properties in Adult Auditory Cortex. Cerebral Cortex, 25(8), 2083–2094. 10.1093/cercor/bhu013

Mowery, T. M., Kotak, V. C., & Sanes, D. H. (2016). The onset of visual experience gates auditory cortex critical periods. Nature Communications, 7(1), Article 1. 10.1038/ncomms10416

Murthy, S., Kane, G. A., Katchur, N. J., Lara Mejia, P. S., Obiofuma, G., Buschman, T. J., McEwen, B. S., & Gould, E. (2019). Perineuronal Nets, Inhibitory Interneurons, and Anxiety-Related Ventral Hippocampal Neuronal Oscillations Are Altered by Early Life Adversity. Biological Psychiatry, 85(12), 1011–1020. 10.1016/j.biopsych.2019.02.021

Nardou, R., Sawyer, E., Song, Y. J., Wilkinson, M., Padovan-Hernandez, Y., de Deus, J. L., Wright, N., Lama, C., Faltin, S., Goff, L. A., Stein-O’Brien, G. L., & Dölen, G. (2023). Psychedelics reopen the social reward learning critical period. Nature, 618(7966), Article 7966. 10.1038/s41586-023-06204-3

Nelson, C. A., Zeanah, C. H., Fox, N. A., Marshall, P. J., Smyke, A. T., & Guthrie, D. (2007). Cognitive Recovery in Socially Deprived Young Children: The Bucharest Early Intervention Project. Science, 318(5858), 1937–1940. 10.1126/science.1143921

Nowicka, D., Soulsby, S., Skangiel-Kramska, J., & Glazewski, S. (2009). Parvalbumin-containing neurons, perineuronal nets and experience-dependent plasticity in murine barrel cortex. European Journal of Neuroscience, 30(11), 2053–2063. 10.1111/j.1460-9568.2009.06996.x

Opendak, M., Gould, E., & Sullivan, R. (2017). Early life adversity during the infant sensitive period for attachment: Programming of behavioral neurobiology of threat processing and social behavior. Developmental Cognitive Neuroscience, 25, 145–159. 10.1016/j.dcn.2017.02.002

Pizzorusso, T., Medini, P., Berardi, N., Chierzi, S., Fawcett, J. W., & Maffei, L. (2002). Reactivation of Ocular Dominance Plasticity in the Adult Visual Cortex. Science, 298(5596), 1248–1251. 10.1126/science.1072699

Poll, S., Mittag, M., Musacchio, F., Justus, L. C., Giovannetti, E. A., Steffen, J., Wagner, J., Zohren, L., Schoch, S., Schmidt, B., Jackson, W. S., Ehninger, D., & Fuhrmann, M. (2020). Memory trace interference impairs recall in a mouse model of Alzheime’s disease. Nature Neuroscience, 23(8), 952–958. 10.1038/s41593-020-0652-4

Polley, D. B., Thompson, J. H., & Guo, W. (2013). Brief hearing loss disrupts binaural integration during two early critical periods of auditory cortex development. Nature Communications, 4(1), Article 1. 10.1038/ncomms3547

Pompeiano, M., & Colonnese, M. T. (2023). cFOS as a biomarker of activity maturation in the hippocampal formation. Frontiers in Neuroscience, 17. https://www.frontiersin.org/articles/10.3389/fnins.2023.929461

Qin, X., Liu, X.-X., Wang, Y., Wang, D., Song, Y., Zou, J.-X., Pan, H.-Q., Zhai, X.-Z., Zhang, Y.-M., Zhang, Y.-B., Hu, P., & Zhang, W.-H. (2021). Early life stress induces anxiety-like behavior during adulthood through dysregulation of neuronal plasticity in the basolateral amygdala. Life Sciences, 285, 119959. 10.1016/j.lfs.2021.119959

Raineki, C., Opendak, M., Sarro, E., Showler, A., Bui, K., McEwen, B. S., Wilson, D. A., & Sullivan, R. M. (2019). During infant maltreatment, stress targets hippocampus, but stress with mother present targets amygdala and social behavior. Proceedings of the National Academy of Sciences, 116(45), 22821–22832. 10.1073/pnas.1907170116

Ramsaran, A. I., Schlichting, M. L., & Frankland, P. W. (2019). The ontogeny of memory persistence and specificity. Developmental Cognitive Neuroscience, 36, 100591. 10.1016/j.dcn.2018.09.002

Ramsaran, A. I., Wang, Y., Golbabaei, A., Aleshin, S., de Snoo, M. L., Yeung, B. A., Rashid, A. J., Awasthi, A., Lau, J., Tran, L. M., Ko, S. Y., Abegg, A., Duan, L. C., McKenzie, C., Gallucci, J., Ahmed, M., Kaushik, R., Dityatev, A., Josselyn, S. A., & Frankland, P. W. (2023). A shift in the mechanisms controlling hippocampal engram formation during brain maturation. Science, 380(6644), 543–551. 10.1126/science.ade6530

Rao-Ruiz, P., Yu, J., Kushner, S. A., & Josselyn, S. A. (2019). Neuronal competition: Microcircuit mechanisms define the sparsity of the engram. Current Opinion in Neurobiology, 54, 163–170. 10.1016/j.conb.2018.10.013

Rashid, A. J., Yan, C., Mercaldo, V., Hsiang, H.-L. (Liz), Park, S., Cole, C. J., De Cristofaro, A., Yu, J., Ramakrishnan, C., Lee, S. Y., Deisseroth, K., Frankland, P. W., & Josselyn, S. A. (2016). Competition between engrams influences fear memory formation and recall. Science, 353(6297), 383–387. 10.1126/science.aaf0594

Rubin, D. C., & Schulkind, M. D. (1997). The distribution of autobiographical memories across the lifespan. Memory & Cognition, 25(6), 859–866. 10.3758/BF03211330

Ruediger, S., Vittori, C., Bednarek, E., Genoud, C., Strata, P., Sacchetti, B., & Caroni, P. (2011). Learning-related feedforward inhibitory connectivity growth required for memory precision. Nature, 473(7348), Article 7348. 10.1038/nature09946

Sale, A., Putignano, E., Cancedda, L., Landi, S., Cirulli, F., Berardi, N., & Maffei, L. (2004). Enriched environment and acceleration of visual system development. Neuropharmacology, 47(5), 649–660. 10.1016/j.neuropharm.2004.07.008

Scott, J. P. (1962). Critical Periods in Behavioral Development. Science, 138(3544), 949–958. 10.1126/science.138.3544.949

Smith, S. L., & Trachtenberg, J. T. (2007). Experience-dependent binocular competition in the visual cortex begins at eye opening. Nature Neuroscience, 10(3), Article 3. 10.1038/nn1844

Sztainberg, Y., & Chen, A. (2010). An environmental enrichment model for mice. Nature Protocols, 5(9), Article 9. 10.1038/nprot.2010.114

Tan, L., Tring, E., Ringach, D. L., Zipursky, S. L., & Trachtenberg, J. T. (2020). Vision Changes the Cellular Composition of Binocular Circuitry during the Critical Period. Neuron, 108(4), 735–747.e6. 10.1016/j.neuron.2020.09.022

Tanti, A., Belliveau, C., Nagy, C., Maitra, M., Denux, F., Perlman, K., Chen, F., Mpai, R., Canonne, C., Théberge, S., McFarquhar, A., Davoli, M. A., Belzung, C., Turecki, G., & Mechawar, N. (2022). Child abuse associates with increased recruitment of perineuronal nets in the ventromedial prefrontal cortex: A possible implication of oligodendrocyte progenitor cells. Molecular Psychiatry, 27(3), Article 3. 10.1038/s41380-021-01372-y

Tooley, U. A., Bassett, D. S., & Mackey, A. P. (2021). Environmental influences on the pace of brain development. Nature Reviews Neuroscience, 22(6), Article 6. 10.1038/s41583-021-00457-5

Travaglia, A., Bisaz, R., Sweet, E. S., Blitzer, R. D., & Alberini, C. M. (2016). Infantile amnesia reflects a developmental critical period for hippocampal learning. Nature Neuroscience, 19(9), Article 9. 10.1038/nn.4348

Trouche, S., Sasaki, J. M., Tu, T., & Reijmers, L. G. (2013). Fear Extinction Causes Target-Specific Remodeling of Perisomatic Inhibitory Synapses. Neuron, 80(4), 1054–1065. 10.1016/j.neuron.2013.07.047

Vetere, G., Xia, F., Ramsaran, A. I., Tran, L. M., Josselyn, S. A., & Frankland, P. W. (2021). An inhibitory hippocampal–thalamic pathway modulates remote memory retrieval. Nature Neuroscience, 24(5), Article 5. 10.1038/s41593-021-00819-3

Villers-Sidani, E. de, Chang, E. F., Bao, S., & Merzenich, M. M. (2007). Critical Period Window for Spectral Tuning Defined in the Primary Auditory Cortex (A1) in the Rat. Journal of Neuroscience, 27(1), 180–189. 10.1523/JNEUROSCI.3227-06.2007

Wang, D., & Fawcett, J. (2012). The perineuronal net and the control of CNS plasticity. Cell and Tissue Research, 349(1), 147–160. 10.1007/s00441-012-1375-y

Wang, X.-D., Labermaier, C., Holsboer, F., Wurst, W., Deussing, J. M., Müller, M. B., & Schmidt, M. V. (2012). Early-life stress-induced anxiety-related behavior in adult mice partially requires forebrain corticotropin-releasing hormone receptor 1. European Journal of Neuroscience, 36(3), 2360–2367. 10.1111/j.1460-9568.2012.08148.x

Willis, A., Pratt, J. A., & Morris, B. J. (2021). BDNF and JNK Signaling Modulate Cortical Interneuron and Perineuronal Net Development: Implications for Schizophrenia-Linked 16p11.2 Duplication Syndrome. Schizophrenia Bulletin, 47(3), 812–826. 10.1093/schbul/sbaa139

Wills, T.J., Cacucci, F., Burgees, N., & Keefe, J. (2010). Development of the Hippocampal Cognitive Map in Preweanling Rats. Science, 328(5985), 1573 1576. 10.1126/science.1188224

Woodcock, E. A., & Richardson, R. (2000). Effects of environmental enrichment on rate of contextual processing and discriminative ability in adult rats. Neurobiology of Learning and Memory, 73(1), 1–10. 10.1006/nlme.1999.3911

Xue, J., Brawner, A. T., Thompson, J. R., Yelhekar, T. D., Newmaster, K. T., Qiu, Q., Cooper, Y. A., Yu, C. R., Ahmed-Braima, Y. H., Kim, Y., & Lin, Y. (2024). Spatiotemporal Mapping and Molecular Basis of Whole-brain Circuit Maturation (p. 2024.01.03.572456). bioRxiv. 10.1101/2024.01.03.572456

Ye, Q., & Miao, Q. (2013). Experience-dependent development of perineuronal nets and chondroitin sulfate proteoglycan receptors in mouse visual cortex. Matrix Biology, 32(6), 352–363. 10.1016/j.matbio.2013.04.001

Zhang, L. I., Bao, S., & Merzenich, M. M. (2002). Disruption of primary auditory cortex by synchronous auditory inputs during a critical period. Proceedings of the National Academy of Sciences, 99(4), 2309–2314. 10.1073/pnas.261707398

